# Neural-circuit basis of song preference learning in fruit flies

**DOI:** 10.1101/2023.10.24.563693

**Authors:** Keisuke Imoto, Yuki Ishikawa, Yoshinori Aso, Jan Funke, Ryoya Tanaka, Azusa Kamikouchi

## Abstract

As observed in human language learning and song learning in birds, the fruit fly *Drosophila melanogaster* changes its’ auditory behaviors according to prior sound experiences. Female flies that have heard male courtship songs of the same species are less responsive to courtship songs of different species. This phenomenon, known as song preference learning in flies, requires GABAergic input to pC1 neurons in the central brain, with these neurons playing a key role in mating behavior by integrating multimodal sensory and internal information. The neural circuit basis of this GABAergic input, however, has not yet been identified.

Here, we find that pCd-2 neurons, totaling four cells per hemibrain and expressing the sex-determination gene *doublesex*, provide the GABAergic input to pC1 neurons for song preference learning. First, RNAi-mediated knockdown of GABA production in pCd-2 neurons abolished song preference learning. Second, pCd-2 neurons directly, and in many cases mutually, connect with pC1 neurons, suggesting the existence of reciprocal circuits between pC1 and pCd-2 neurons. Finally, GABAergic and dopaminergic inputs to pCd-2 neurons are necessary for song preference learning. Together, this study suggests that reciprocal circuits between pC1 and pCd-2 neurons serve as a sensory and internal state-integrated hub, allowing flexible control over female copulation. Consequently, this provides a neural circuit model that underlies experience-dependent auditory plasticity.

**Significance:** To find a suitable mate, an organism must adapt its behavior based on past experiences. In the case of *Drosophila*, female assessments of male song signals, which contain information about the status and species of the sender, are experience dependent. Here, we show that reciprocal circuits in the central brain modulate the female’s song response depending on her previous auditory experiences. These circuits exhibit feedback and lateral inhibition motifs, and are regulated by dopaminergic and GABAergic inputs. While the effects of prior auditory experiences on sound responsiveness have been extensively studied in other species, our research advances the use of *Drosophila* as a model for dissecting the circuitry underlying experience-dependent auditory plasticity at single-cell resolution.

## Introduction

Many animals, ranging from humans to birds to insects, produce sounds to communicate with others of the same species. Each species typically has its own sounds, such as languages, calls, or songs. Accordingly, the auditory capability to discriminate key sound communication features is indispensable (1, 2). Studies of vocal learning in infants or juvenile birds have revealed that such discrimination abilities are shaped by the interaction between nature and nurture, relying on innate abilities and experience-dependent auditory plasticity, respectively (3, 4). Human infants hone their capability to make phonetic distinctions by repeated exposure to their native language (5): The human brain starts becoming attuned to the native language a few days after birth, due to prenatal and/or short-term postnatal exposure to the native language (6, 7). Similarly, juvenile songbirds develop their auditory discrimination ability during song learning by hearing tutor songs in the two months after birth (4). Other animals, such as mice and frogs, also utilize courtship sounds for mating (8, 9). The neural circuit mechanism of how their brain is tuned to their unique communication sound, however, has just started to be identified (10, 11), and the cellular basis of sound experience-dependent learning remains to be elucidated.

As observed in the mating communication of many animals, male fruit flies (*Drosophila melanogaster*) emit various signals to attract females, including the near-field sound known as the courtship song which is produced by wing vibrations (12, 13). The major component of the courtship song of *D. melanogaster* is the pulse song, i.e., a repetition of sound pulses (14). The inter-pulse interval (IPI) of the pulse song differs among sibling *Drosophila* species and significantly affects female mate choice, with increased mating receptivity to conspecific courtship songs rather than heterospecific songs (13, 15). The auditory pathways from sensory neurons to higher-order brain neurons are organized to tune selectively to the conspecific song and thus facilitate mating acceptance in female flies (16–18). Several neurons in this circuit express the sex-specific transcription factor *doublesex* (*dsx*), which plays a role in their sexual differentiation (18–20). In the fly brain, *dsx*-expressing neurons form several clusters (five clusters in both males and females, with four male-specific and one female-specific) (19) and previous studies have reported the importance of *dsx-* expressing neurons in mating behaviors (21). Among them, female pC1 neurons, which innervate the lateral protocerebral complex and superior-medial protocerebrum (SMP) in the brain, serve as a key regulator of female mating behavior by enhancing copulation receptivity when activated (20).

Our previous study showed that both male and female fruit flies acquire a preference for songs with conspecific IPIs over songs with heterospecific IPIs following prior exposure to the conspecific song (22). Further studies in females revealed the importance of the neurotransmitter γ-aminobutyric acid (GABA), as GABAergic inputs to pC1 neurons are necessary for song preference learning (22). Studies on the equivalent neuronal mechanism in songbirds have also suggested GABA to be a key regulator in forming auditory memories during the song-learning process: some higher-order auditory cortical neurons become tuned to the tutor song through the recruitment of GABAergic inhibition when exposed in early life (11). However, the neural circuit basis of how sound exposure tunes the higher-order integration center that controls the auditory responses remains unclear in either case.

To address this issue, we used a fruit-fly model to identify GABAergic neurons within neural circuits that mediate song preference learning. First, using an intersectional strategy we showed that pCd-2 neurons, a small number of GABAergic neurons that express *dsx*, are the responsible neurons for song preference learning. Second, mining of the *Drosophila* hemibrain connectome database revealed synaptic connections between pCd-2 neurons and pC1 neurons, including reciprocal ones. Finally, we showed that pCd-2 neurons receive GABAergic and dopaminergic inputs via GABA_A_ receptors (Rdl) and Dop1R2 receptors, respectively, that contribute differently to song preference learning. Taken together, this study reveals at a single-cell resolution the fundamental neural circuit that underlies song preference learning in fruit flies.

## Results

### *doublesex*-expressing GABAergic neurons control song preference learning

Approximately 6,000 GABAergic neurons are present in the adult fly brain (23). To identify GABAergic neurons involved in song preference learning, we focused on the *dsx*-expressing neurons (approximately 66 and 315 neurons in the brain and ventral nerve cord of females, respectively) (24), which are involved in female mating receptivity. To genetically label *dsx*-expressing GABAergic neurons, we utilized an intersectional strategy to generate a *dsx*∩*Gad1* driver, in which *Gad1* encodes the rate-limiting enzyme in the GABA synthesis pathway (Fig. 1 *A* and *B*; see Materials and Methods for fly strains). We found four *dsx*-expressing GABAergic neurons in the hemibrain, and approximately 200 such neurons in the ventral nerve cord (VNC) in females, consistent with numbers reported in previous studies (Fig. 1*A*) (24).

**Fig. 1.**
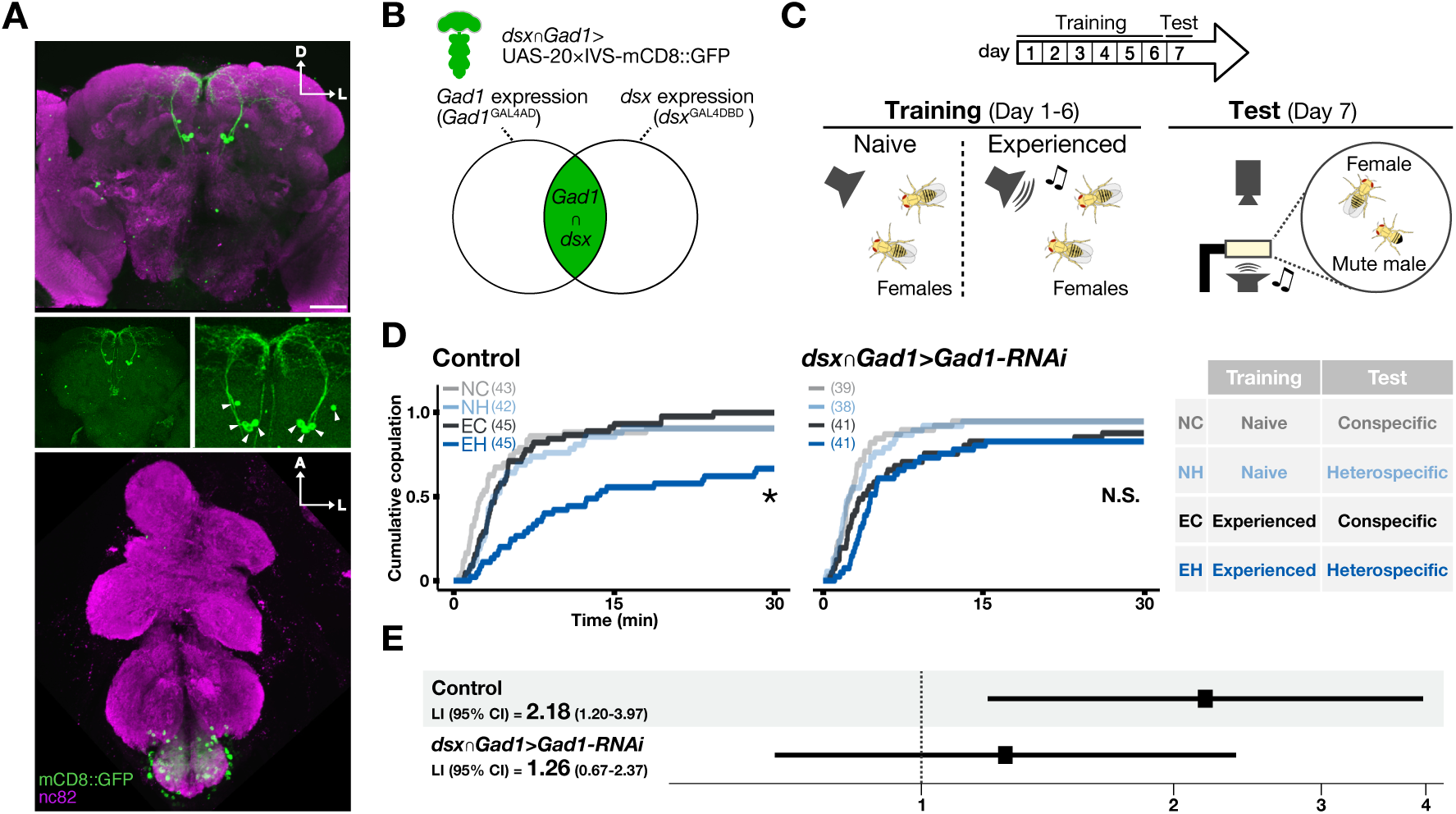
*doublesex* expressing GABAergic neurons are necessary for song preference learning. (A) Expression pattern of *dsx*∩*Gad1* driver in the brain (top and middle-left; anterior view) and ventral nerve chord (bottom; ventral view) in females. A magnified view of the cell bodies (arrowheads) is shown in the middle-right. Scale bars, 50 μm. A, anterior; D, dorsal; L, Lateral (the same in the following figures). (B) Venn diagram of the genetic intersection. The population labeled by the *dsx*∩*Gad1* driver is shown in green. (C) Experimental scheme for song preference learning. In the training session, females in the experienced condition are exposed to the conspecific song for the first 6 days after eclosion while naïve females are kept in silence. During the test session on the 7^th^ day, females of both conditions are paired with mute males and exposed to either conspecific or heterospecific song. (D) Song preference learning in *Gad1* knockdown females. The number of trials for each group is shown in parentheses. NC, naïve flies tested with the conspecific song; NH, naïve flies tested with heterospecific song.; EC, experienced flies tested with the conspecific song; EH, experienced flies tested with heterospecific song. Not significant (N.S.), p>0.05; *, p<0.05; log-logistic AFT model. (E) The learning index (LI) estimated based on the cumulative copulation rate using log-logistic AFT model. The horizontal axis uses a natural logarithm scale (See *SI Appendix,* Supporting Information Text for details). The squares indicate estimated LI and horizontal lines indicate 95% confidence intervals (CIs) (same in the following figures).

Next, we examined whether *dsx*-expressing GABAergic neurons contribute to song preference learning. The song preference learning paradigm comprised a training session and a subsequent test session (Fig. 1*C*) (22). During the training session, female flies in the experienced condition were exposed to an artificial conspecific song, while those in the naive condition were kept without sound exposure. During the test session, female flies in both conditions were paired with mute males and exposed to one of two artificial songs (conspecific or heterospecific song). In this test, the cumulative copulation rate serves as a readout of female receptivity for the courting male (Fig. 1*D*; see also Materials and Methods). To assess the effect of song exposure, we defined the song preference learning index, LI, based on the accelerated failure time (AFT) model (25, 26). In brief, the LI value represents the magnitude of learning: when the LI and its confidence intervals (CIs) deviate from 1, the flies are detected as showing song preference learning (See Materials and Methods; LI of wild-type flies shown in *SI Appendix,* Fig. *S1*).

Using the *dsx*∩*Gad1* driver, we suppressed *Gad1* expression specifically in *dsx*-expressing GABAergic neurons (see Materials and Methods for fly strains). Control flies without any knockdown showed song preference learning: naïve flies rapidly initiated copulation behaviors after being exposed to either the conspecific or heterospecific song (NC and NH, respectively), whereas experienced flies showed a lower copulation rate during exposure to the heterospecific song, but not the conspecific song (EH and EC, respectively) (Fig. 1*D*, left). The AFT model detected a significant interaction between song experience and test song type, indicating that control flies showed experience-dependent reduction of female receptivity to the heterospecific song (Fig. 1*E*; LI = 2.18; 95% CI, 1.20 to 3.97; p = 0.011; log-logistic AFT model; see Materials and Methods). In contrast, *Gad1* knockdown females lost the experience-dependent reduction of copulation (Fig. 1*D*, right): they showed a high copulation rate under all conditions, and no significant interaction was detected (Fig. 1*D*, right and 1E; LI = 1.26; 95% CI, 0.67 to 2.37; p = 0.48; log-logistic AFT model). These results show that suppression of GABA synthesis in *dsx*-expressing GABAergic neurons abolished song preference learning, indicating that neuronal signals from *dsx*-expressing GABAergic neurons are involved in song preference learning.

### pCd-2 neurons in the brain are the key GABAergic neurons for song preference learning

*dsx*-expressing GABAergic neurons are found in the brain and VNC (Fig. 1*A*). To test whether the brain subset contains the neurons responsible for song preference learning, we generated a *dsx*∩*Gad1*^brain^ driver strain by combining the *dsx*∩*Gad1* strain with an *Otd-FLP*, *tubP FRT-Gal80-FRT* strain. This driver induces GAL4-dependent gene expression only in the brain in principle, since *Otd-FLP* drives FLP expression specifically in the central brain (27). The *dsx*∩*Gad1*^brain^ clearly labeled the neural subset in the brain but only limited labeling was observed in the VNC (N = 4; Fig. 2 *A-C*). All labeled neurons in the brain, 4 neurons per hemibrain, have cell bodies lying at the posterior-dorsal side of the brain, ventral to the protocerebral bridge, and extend neurites to the superior medial protocerebrum (SMP) and ipsilateral gnathal ganglia (GNG) (Fig. 2 *B* and *C*). Their number, locations in the brain, and neurite morphology are virtually identical to those observed in the *dsx*∩*Gad1* driver (Fig. 1*A*), indicating that these two intersections labeled the same set of neurons in the brain.

**Figure 2.**
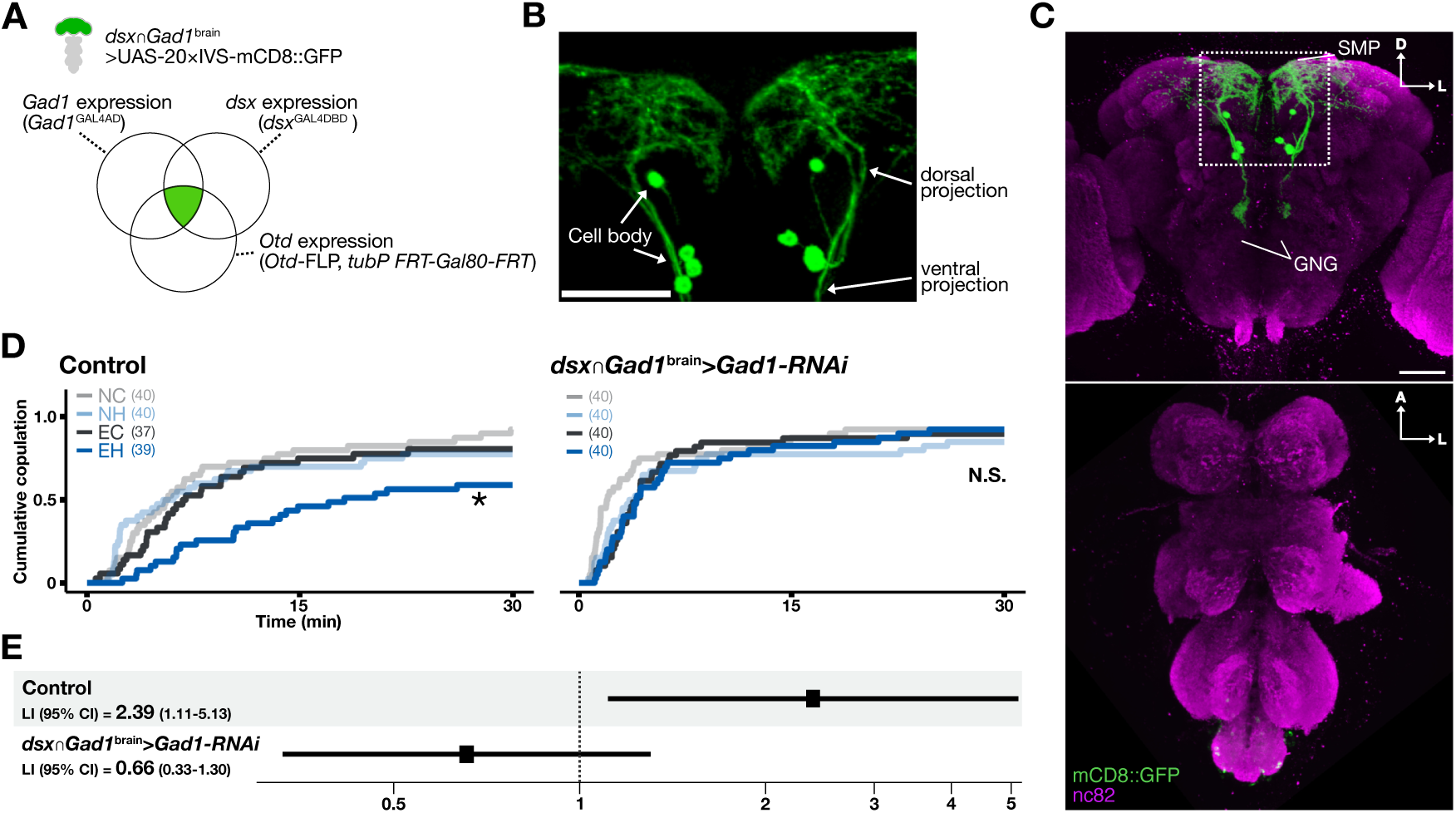
pCd-2 neurons control song preference learning. (A) Venn diagram of genetic intersection. The area containing *dsx*∩*Gad1*^brain^ -labeled neurons is shown in green. (B-C) Expression pattern of *dsx*∩*Gad1*^brain^ driver in females. Magnified view of the neurites and cell bodies located in the superior medial protocerebrum (SMP) (B), and overview in the central brain (C, top; anterior view) and ventral nerve cord (C, bottom; ventral view) are shown. GNG, gnathal ganglia. Scale bars, 50 µm. (D) Cumulative copulation rates in control and pCd-2 specific *Gad1* knockdown groups. NC, naïve flies tested with the conspecific song; NH, naïve flies tested with heterospecific song.; EC, experienced flies tested with the conspecific song; EH, experienced flies tested with heterospecific song. The number of trials in each group is shown in parentheses. N.S., not significant; *p<0.05; log-logistic AFT model. (E) Learning index (LI) with 95% confidence interval (CI).

To examine whether *dsx*-expressing GABAergic neurons in the brain are involved in song preference learning, we knocked down *Gad1* expression in these cells using the *dsx*∩*Gad1*^brain^ driver. Control females exhibited song preference learning as previously shown for wild-type females (*SI Appendix,* Fig. S1), and as such females showed a reduced copulation rate in response to the heterospecific song under the experienced training condition (Fig. 2*D*, left; Fig. 2*E*; LI = 2.39; 95% CI, 1.11 to 5.13; p= 0.026; log-logistic AFT model). In contrast, females with *Gad1*-knockdown in *dsx*∩*Gad1*^brain^ neurons showed high copulation rates in response to both conspecific and heterospecific songs regardless of naïve and experienced training condition, demonstrating no song preference learning (Fig. 2*D*, right and 2*E*; LI = 0.66; 95% CI, 0.33 to 1.30; p= 0.23; log-logistic AFT model; also see *SI Appendix,* Fig. S2 for another RNAi strain). We also performed a genetic intersection with the *tsh-Gal80* strain, which suppresses expression in most VNC neurons (28). This intersection, which is hereafter referred to as *dsx*∩*Gad1*^brain2^, clearly labeled the neural subset in the brain but showed only limited labeling in the VNC (N = 4; *SI Appendix,* Fig. S3 *A* and *B*). *Gad1* knockdown by the *dsx*∩*Gad1*^brain2^ driver disrupted song preference learning (*SI Appendix,* Fig. S3 *C* and D). As a result of finding disruption in song preference learning in *Gad1* knockdown flies from two independent “brain-dominant” strains, we concluded that the four *dsx*-expressing GABAergic neurons in the brain are the GABAergic neurons responsible for song preference learning.

Previous studies identified seven to eight clusters of *dsx*-expressing neurons in the female brain, i.e., pC1, pC2l, pC2m, pCd-1, pCd-2, pMN1, pMN2, and aDN clusters, with distinct cell-body locations and neurite morphologies (18, 19, 29). The number, cell-body locations, and neurite morphologies of *dsx*∩*Gad1*^brain^ -labeled neurons resemble those of pCd-2 neurons and are distinct from other *dsx*-expressing neuronal clusters (Fig. 2*B*; See *SI Appendix,* Supporting Information Text for detailed morphology) (19, 29), indicating that *dsx*∩*Gad1*^brain^ neurons belong to the pCd-2 cluster. Taken together, we concluded that pCd-2 neurons control song preference learning through GABAergic signaling pathways.

### pCd-2 neurons connect to pC1 neurons

pCd-2 neurites are reported to spatially overlap with pC1 neurites in the SMP (19). Given that GABAergic input to pC1 neurons regulates song preference learning (22), we speculated that pCd-2 neurons might directly connect to pC1 neurons to regulate song preference learning. To assess the synaptic connections, we performed GFP reconstitution across synaptic partners (GRASP) analysis (22) by expressing spGFP_1-10_ and spGFP_11_ in pCd-2 neurons and pC1 neurons, respectively. In all cases tested (N=4), reconstructed GFP signals (GRASP signals) were detected at the posterior side of the SMP (Fig. 3*A*). It is thus highly likely that pCd-2 neurons have direct synaptic connections with pC1 neurons.

**Figure 3.**
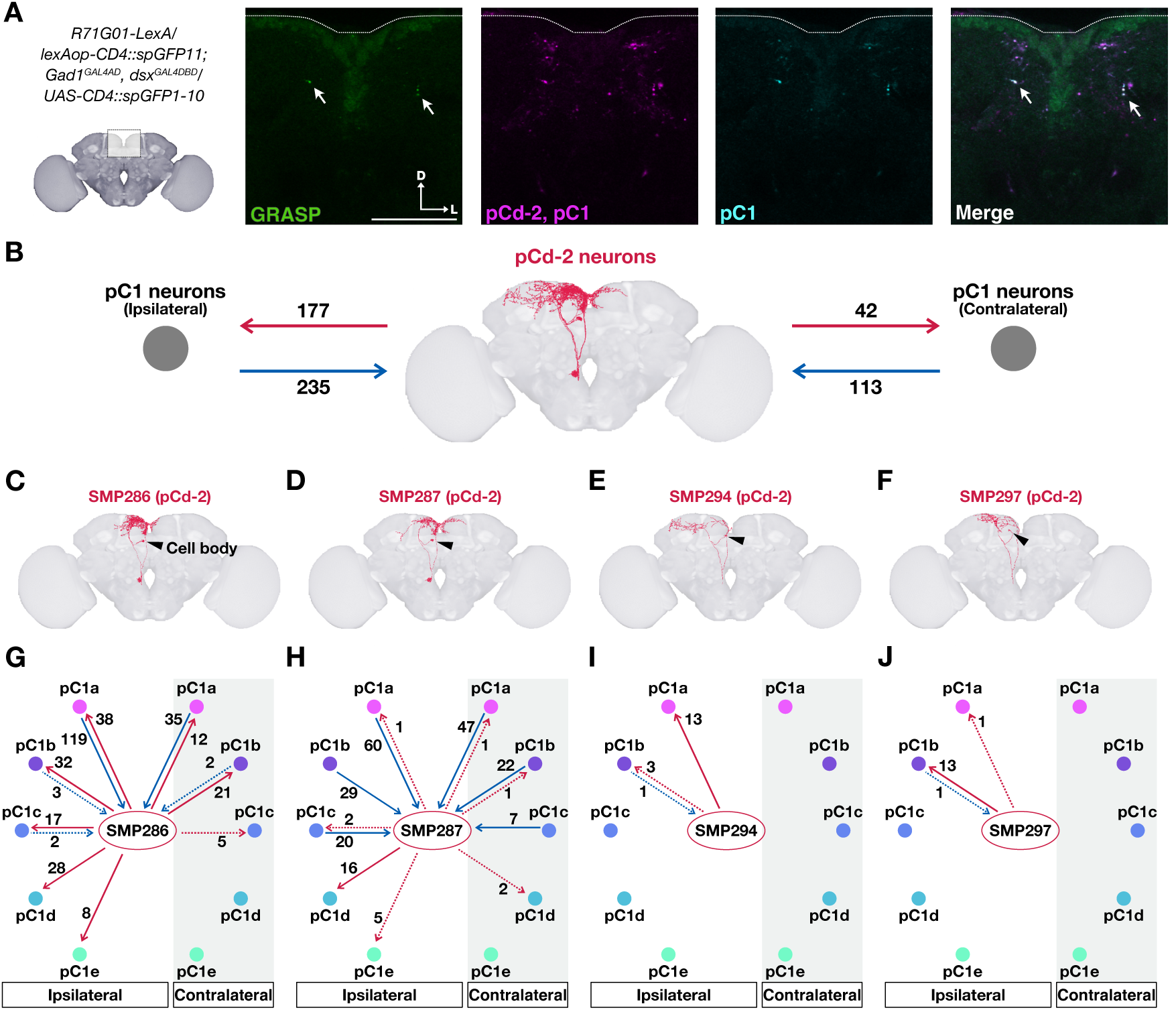
Synaptic connections between pC2d and pC1. (A) GRASP between pCd-2 neurons and pC1 neurons. The brain region shown in the middle to right panels is outlined in the left panel. GRASP signals (green), spGFP_1-10_ and spGFP_11_ expression in pCd-2 neurons and pC1 neurons respectively (magenta), spGFP_11_ expression in pC1 neurons (light blue), and merged image are shown (See *SI Appendix,* Table S1 for the genotype). Scale bar, 50 μm. (B) Synaptic connections between pC1 neurons and pCd-2 neurons. Red and blue arrows depict the output and input synapses of pCd-2 neurons to/from pC1 neurons, respectively. Numbers on arrows indicate the number of synapses. (C-F) Single pCd-2 neurons in the right hemibrain of the FlyEM dataset. (G-J) Synaptic connections between pCd-2 neurons and pC1 neurons based on the FlyEM dataset. Numbers on arrows indicate the number of synapses. Weak connections (fewer than 6 synapses) (18) are shown in dotted arrows.

To reveal the synaptic connections between pC1 and pCd-2 neurons at the single cell level, we analyzed the FlyEM dataset, an EM-based connectome dataset of an adult female fly brain (hemibrain: v1.2.1) (30). pC1 neurons are comprised of five cells, namely, pC1a, pC1b, pC1c, pC1d, and pC1e (31) (see *SI Appendix,* Table S2 for the FlyEM neuron IDs). These subtypes have been suggested to control different types of female behaviors, i.e., aggression and copulation receptivity (18)). To identify pCd-2 neurons labeled by the *dsx∩Gad1^brain^* driver in the FlyEM dataset, we searched for neurons whose morphologies resemble those of pCd2 neurons and have GABA synapses to pC1 neurons (threshold = 10 synapses; *SI Appendix*, Supporting Information Text) using a neurotransmitter classification algorithm (32). We identified four neurons in a hemibrain, named SMP286, SMP287, SMP294, and SMP297, which share comparable cell body locations and projection patterns with pCd-2 neurons (Fig. 3 *B*-*F*; FlyEM neuron IDs are listed in *SI Appendix,* Table S2). The distributions of postsynaptic sites in these four neurons, mapped in the FlyEM, were consistent with the pCd-2 dendritic regions labeled by a postsynaptic marker DenMark (33) driven by the *dsx*∩*Gad1*^brain^ driver (*SI Appendix,* Fig. S4). Thus, we classified these four neurons as pCd-2 neurons, GABAergic neurons projecting to pC1 neurons.

By investigating synaptic connections between these four pCd-2 neurons and pC1 neurons with the FlyEM dataset (Fig. 3*B*), we classified pCd-2 neurons into two types. One is the intensive connection type (SMP286 and SMP287), which has many synapses with several types of pC1 neurons (Fig. 3 *G* and *H*). The other is the sparse connection type (SMP294 and SMP297), with fewer synapses with a subset of pC1 neurons (Fig. 3 *I* and *J*). Among the intensive connection-type pCd-2 neurons, SMP286 has many bilateral output synapses to pC1 neurons, while the number and variety of ipsilateral outputs exceed that of contralateral outputs. Importantly, SMP286 receives intensive synaptic inputs from pC1a neurons of both brain sides, indicating reciprocal synaptic connections between SMP286 and bilateral pC1a neurons. The other intensive connection-type pCd-2 neuron, SMP287, has output synapses mainly to the ipsilateral pC1d neuron but receives many synaptic inputs bilaterally from pC1a/b/c neurons. In contrast, the sparse connection-type pCd-2 neurons have output synapses almost exclusively to pC1 (Fig. 3 *I* and *J*). pC1a and pC1b receive inputs from ipsilateral SMP294 and SMP297, respectively, suggesting a specific regulation from these pCd-2 neurons to a corresponding pC1 neuron. Taken together, GRASP and the FlyEM connectome analyses revealed the synaptic connectivity between pC1 and pCd-2 neurons. Interestingly, there are direct synaptic connections from SMP294 to SMP286 and SMP286 to SMP287 but not vice versa, suggesting a unidirectional information flow among these three pCd-2 neurons (*SI Appendix,* Fig. S6A, Table S3). It is possible that all types of pCd-2 neurons are involved in the GABAergic control of pC1 neurons, while the specific combination of target pC1 neurons and synaptic weights differ between each type of pCd-2 neuron (Fig. 3 *G-J* and *SI Appendix,* Fig. S5). Furthermore, two of the pCd-2 neurons (SMP286 and SMP287) receive many direct synaptic inputs from pC1 neurons, suggesting the existence of reciprocal connectivity between these pCd-2 neurons and pC1 neurons that together exhibit feedback and lateral inhibition motifs.

FlyEM database mining also revealed that two intensive-type pCd-2 neurons have further direct synaptic connections with other neurons involved in female copulation receptivity (*SI Appendix,* Fig. S6B). SMP286 and SMP287 receive synaptic inputs from SAG neurons, activation of which is known to depolarize pC1 neurons and increases female receptivity (31, 34, 35). SMP286 sends synaptic output to vpoDN, which controls vaginal plate opening to accept copulation (*SI Appendix,* Table S3) (35). vpoDN also receives direct synaptic input from pC1 neurons (35), suggesting a feed-forward circuit motif from SAG-neurons to vpoDN neurons through pC1-pCd-2 reciprocal circuits (*SI Appendix,* Fig. S6). These findings suggest that two intensive-type pCd-2 neurons are embedded in the neural circuit comprised of SAG, pC1, and vpoDN, possibly to modulate female receptivity and copulation acceptance.

### GABA and dopamine signaling to pCd-2 neurons

Our findings suggest that pCd-2 neurons send GABAergic signals to pC1 neurons to suppress female receptivity in response to heterospecific song exposure. To further explore how pCd-2 activity is modulated by upstream neurons, we screened neurotransmitter receptors expressed in pCd-2 neurons using a single-cell transcriptome of adult fly brains (23, 36). To focus on the gene expression profiles of pCd-2 neurons, we selected cells which express *Gad1*, *dsx,* and *elav* genes, among which *elav* serves as a pan-neuronal marker (37). Many neurons in this population expressed several types of receptors, including GABA_A_-type receptor (Rdl), dopamine 1-like receptor 1 (Dop1R1), dopamine 1-like receptor 2 (Dop1R2), and/or dopamine 2-like receptor (Dop2R) (*SI Appendix*, Fig. S7).

To investigate whether and how these receptors in pCd-2 neurons contribute to song preference learning, we examined the effect of knockdown of each receptor using the *dsx*∩*Gad1*^brain^ driver. Knockdown of either *Dop1R1* or *Dop2R* did not abolish song preference learning, suggesting no or limited involvement in song preference learning (*SI Appendix*, Fig. S8). In contrast, *Rdl* knockdown females lost the learning phenotype (Fig. 4 *A* and *B*; *Rdl* knockdown group: LI = 0.80; 95% CI, 0.42 to 1.51; p= 0.49; control group: LI = 2.39; 95% CI, 1.11 to 5.13; p= 0.026; log-logistic AFT model). The phenotypic changes observed following *Rdl* knockdown in *dsx*∩*Gad1*^brain^ neurons resembled that of *Gad1* knockdown, with experienced females showing as high a copulation rate to the heterospecific song as to the conspecific song (Fig. 4 *A* and *B* vs Fig. 2 *D* and *E*). Interestingly, flies with *Dop1R2* knockdown in *dsx*∩*Gad1*^brain^ neurons exhibited a different type of learning disruption (Fig. 4 *C* and *D*). In the experienced condition, *Dop1R2* knockdown females exhibited lower copulation rates to both the conspecific and heterospecific songs than naïve flies, which led to the loss of the learning phenotype (Fig. 4 *C* and *D*; *Dop1R2* knockdown group: LI = 0.70; 95% CI, 0.33 to 1.49; p= 0.35; control group: LI = 2.30; 95% CI, 1.06 to 4.99; p= 0.039; log-logistic AFT model; see *SI Appendix*, Supporting Information Text for the detailed phenotype). We used restricted mean time lost (RMTL) as an indicator of the cumulative copulation rate to evaluate this change, where a larger RMTL reflects higher copulation acceptance (Fig. 4*E*; See Materials and Methods for details) (15). We found that the response of experienced females to the conspecific song, but not to the heterospecific song, was significantly reduced in the *Dop1R2* knockdown group when compared to that of the control group (conspecific song: p = 1.62e^-12^; heterospecific song: p = 0.32; restricted mean survival time adjusted by Bonferroni). This phenotype contrasted with that in the *Rdl*-knockdown females, which lost the experience-dependent reduction in response to the heterospecific song (Fig. 4*E*; conspecific song: p= 0.057; heterospecific song: p= 0.0022; restricted mean survival time adjusted by Bonferroni correction). These findings suggest that GABAergic and dopaminergic inputs to pCd-2 neurons, through Rdl and Dop1R2 receptors respectively, play different modulatory roles to control song preference learning through pC1 neurons: GABAergic input to pCd2 *via* Rdl receptors is necessary to suppress copulation behavior in response to heterospecific song exposure, and dopaminergic input through Dop1R2 receptors is necessary to facilitate behavioral responses to the conspecific song in experienced flies. While investigating synaptic connections using the FlyEM dataset, we discovered that some pCd-2 neurons (SMP286 and SMP287) receive synaptic input from other pCd-2 neurons, as well as a dopaminergic neuron cluster known as PAL (SI Appendix, Fig. S6A, S6C, Table S3). GABAergic and dopaminergic modulations of these pCd-2 neurons may thus be mediated via signaling from other pCd-2 neurons and PAL neurons, respectively.

**Fig. 4.**
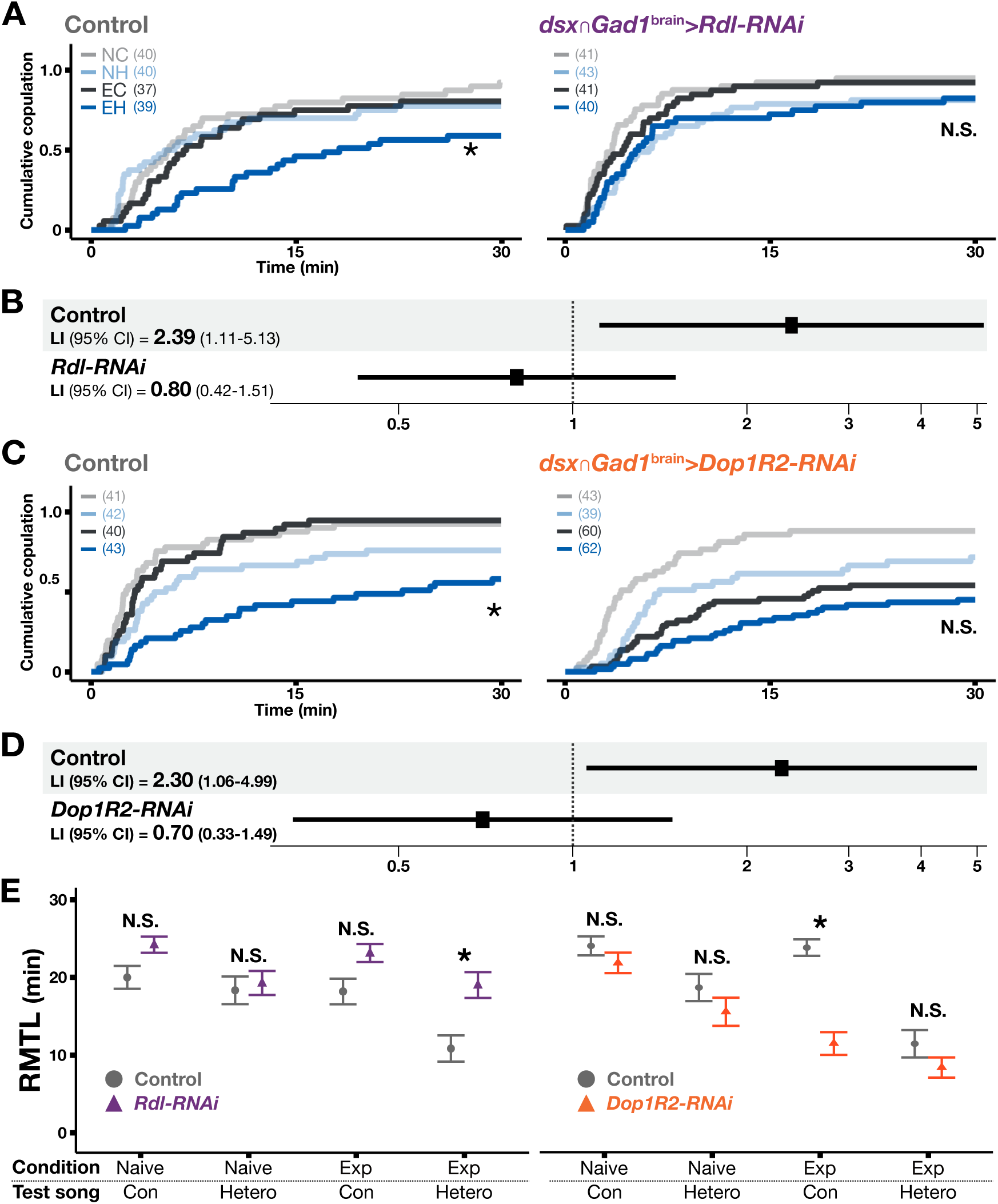
Involvement of GABA and dopamine receptors in song preference learning. (A) Cumulative copulation rates in control and *Rdl* knockdown groups. NC, naïve flies tested with the conspecific song; NH, naïve flies tested with heterospecific song.; EC, experienced flies tested with the conspecific song; EH, experienced flies tested with heterospecific song. The number of trials in each group is shown in parentheses. (B) Learning index (LI) in control and *Rdl* knockdown groups. (C) Cumulative copulation rates in control and *Dop1R2* knockdown groups. (D) LI in control and *Dop1R2* knockdown groups. (E) Restricted mean time lost (RMTL) of cumulative copulation rate for each group. Plots display the average (circle or triangular dot in each box) and standard errors (horizontal bars). Exp, experienced; Con, conspecific song; Hetero, heterospecific song. Log-logistic AFT model (A-D) and restricted mean survival time with Bonferroni correction (E) were used. N.S., not significant; *p<0.05.

## Discussion

Here, we demonstrate that *dsx*-expressing GABAergic pCd-2 neurons are responsible for song preference learning in female fruit flies. pCd-2 neurons have direct connections with pC1 neurons, which regulate female sexual receptivity. A functional analysis of neurotransmitter receptors suggested that GABAergic and dopaminergic inputs to pCd-2 neurons are involved in this learning. These findings refine the neural circuit model that contributes to song response behaviors at the single-cell resolution level (Fig. 5). In this model, song responses of females are regulated by an interaction between innate and experience-dependent pathways (Fig. 5*A*) (22). The innate pathway starts with auditory sensory neurons, which transmit song information finally to pC1 neurons to regulate female receptivity. The experience-dependent pathway modifies the innate pathway through GABAergic pCd-2 neurons which make reciprocal connections with pC-1 neurons (Fig. 5*B*). GABAergic pCd-2 neurons receive GABAergic input through GABA_A_ receptors and dopaminergic input mediated by Dop1R2 receptors. GABAergic signaling to pCd-2 plays a key role in suppressing mating receptivity to the heterospecific song in the experienced flies, while dopamine signals facilitate, and thus maintain, the receptivity for the conspecific song after the experience (Fig. 5*B*). It implies that pCd-2 neurons gate the song response of female flies based on prior sound experiences. One notable feature of the behavioral phenotype of song preference learning in wild-type flies is that the experience affects only the heterospecific song response. This specificity might be achieved via separate GABAergic and dopaminergic modulation pathways to pCd-2 neurons, and its underpinning mechanism should be explored in the future.

**Figure 5.**
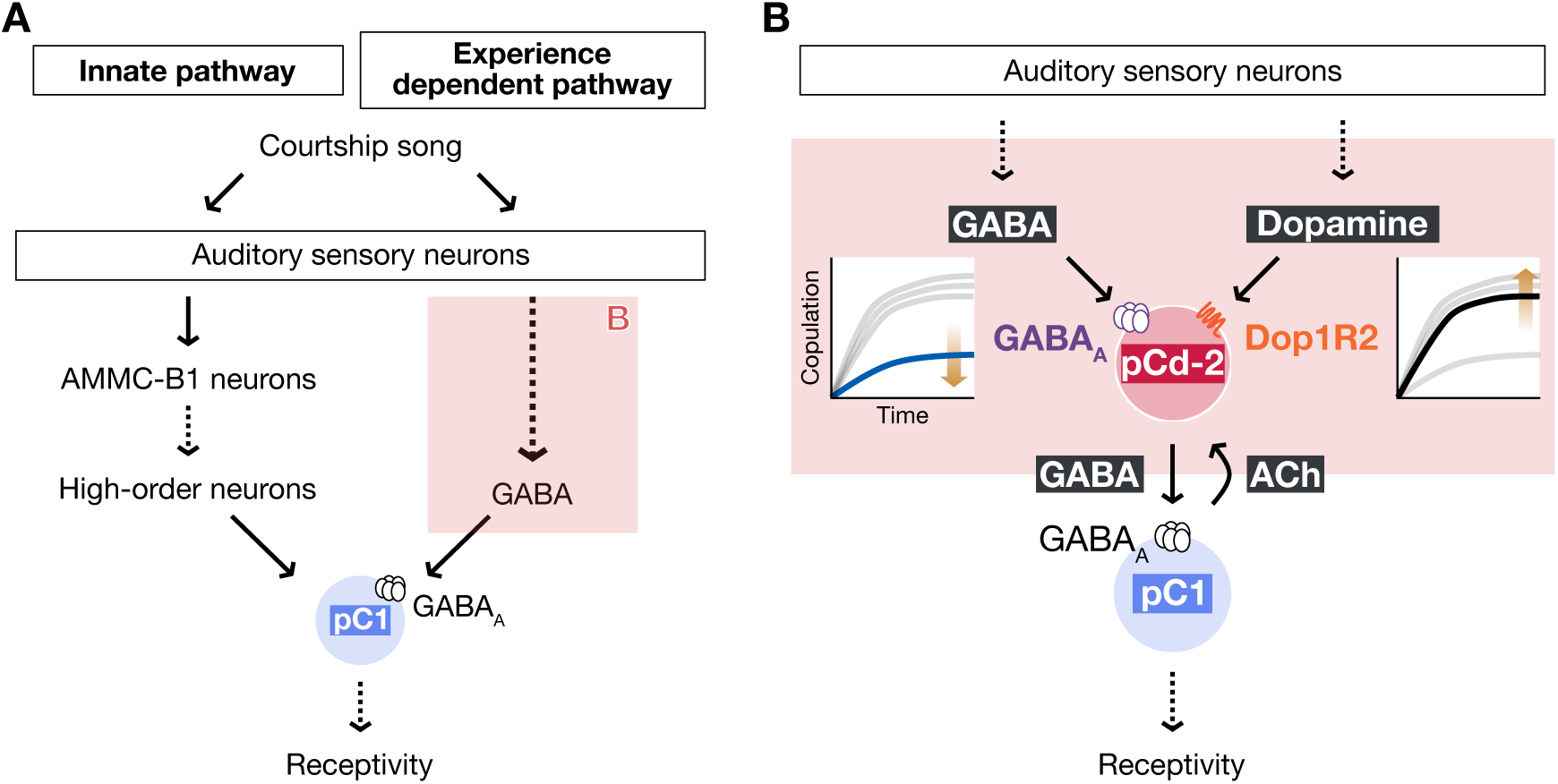
Neural circuit model for song preference learning in flies. (A) A model for experience-dependent tuning of song responses in fruit flies. Flies fine-tune their innate IPI preference through auditory experiences. Modified from (22) with permission. (B) The neural circuit mechanism of experience-dependent modulation. Song preference learning in flies involves at least two distinct mechanisms, the experience-dependent suppression of the response to the heterospecific song (blue line in the copulation plot) and the maintenance of the response to the conspecific song (black line in the copulation plot) after the experience. These two mechanisms are mediated by GABA and dopamine, respectively, transmitted to pCd-2 neurons. pC1 neurons are excitatory cholinergic neurons transmitting acetylcholine (ACh) (31).

### The neural circuit between pCd-2 / pC1 neurons

In this study, we found that a subset of pCd-2 neurons forms reciprocal circuits with pC1 neurons (Fig. 3 *B, G-J*). The pCd-2 neurons are GABAergic, and therefore likely form inhibitory connections to cholinergic pC1 neurons, which, in turn, establish excitatory connections back to pCd-2 neurons (Fig. 5*B*) (31). The reciprocal circuits between these two neuronal types thus likely provide both feedback and lateral inhibitions to pC1 neurons that act as decision-makers for copulation. These inhibitory motifs are commonly observed in the brains of a wide range of animals. In mammals, for example, feedback inhibition in cortical circuits has been suggested to play a significant role in decision computations (38, 39). Lateral inhibition, on the other hand, can mediate competitive interactions between neurons, often leading to contrast enhancement (40). Accordingly, a possible scenario in flies is that the pCd-2 / pC1 reciprocal circuits adjust the copulation decision-making signal based on prior sound experiences through contrasting activity patterns among pC1 subtypes. Further investigation is necessary to explore how GABAergic and dopaminergic signals to pCd-2 neurons modify the activity profile of these reciprocal circuits, as well as to examine how each of the pCd-2 / pC1 cluster neurons are functionally integrated for copulation decision-making.

pCd-2 neurons observed in this study were found in both males and females (*SI Appendix*, Fig. S9), suggesting that these neurons are shared between the sexes. This observation is consistent with a previous study, which reported that pCd-2 neurons also exist in males with a slight sexual dimorphism in neurite structures (19). Since male flies also exhibit song preference learning (22), pCd-2 neurons may play a key role in song preference learning in both sexes. However, a female specific population of pCd-2 neurons, labeled by a distinct set of hemi-drivers, has been reported: these female-specific pCd-2 neurons control post-mating changes in food preference under the regulation of pC1 neurons (41) and are located downstream of the SAG-pC1 pathway, which alters its mode after copulation (35). It will be interesting to further investigate if and how the pCd-2 / pC1 reciprocal circuits underlying song preference learning in female brains interact with the SAG-pC1-pCd-2 pathway involved in post-mating food preference.

### Two different pathways shape song preference after a song experience

GABA and dopamine inputs to pCd-2 neurons are mediated by GABA_A_ (Rdl) and Dop1R2 receptors (Fig. 5*B*). Rdl encodes a subunit of ligand-gated chloride channel GABA_A_, while Dop1R2 is a G-protein-coupled receptor that presumably belongs to the D1-like dopamine receptor group (42). It is known that dopamine signaling via Dop1R2 increases intracellular calcium levels and cAMP (43), whereas GABAergic signaling through GABA_A_ receptors containing Rdl subunits induce fast inhibition due to chloride influx (44). Such distinct signaling cascades in pCd-2 neurons may contribute to the different responses to two song types in experienced females (Fig. 5*B*), though these signaling properties remain to be characterized and validated in pCd-2 neurons.

A candidate source of GABAergic signals to pCd-2 neurons are the pCd-2 neurons themselves, as our connectome analysis detected synaptic connections within pCd-2 neurons (*SI Appendix*, Fig. S6A, Table S3). In the mammalian brain, interactions between GABAergic neurons contribute to synchronized oscillation of cortical neuron activity which is proposed to facilitate neural plasticity (45, 46). Similarly, interactions between GABAergic neurons in flies can potentially contribute to song-experience-dependent neural plasticity. On the other hand, dopaminergic signals to pCd-2 neurons are possibly derived from PAL neurons (*SI Appendix*, Fig. S6C, Table S3). The PAL cluster, located in the superiorlateral protocerebrum in the fly brain (47), is one of the dopaminergic neuron clusters involved in olfactory learning in both males and females (48). During mating behaviors, a dopamine neuron denoted aSP4 in the PAL cluster in males transmits signals to P1 neurons, a male-specific subset of pC1 neurons, via Dop1R2. While this pathway contributes to promoting male mating drive (49), the involvement of the PAL cluster in female mating motivation is not well understood.

In zebra finches, an interaction between dopamine and GABA was reported in a brain region known as the caudomedial nidopallium (NCM), the higher-level auditory cortex in the avian brain that serves as a possible storage site of tutor song memories (50). Most neurons in the NCM co-express D1 receptor (D1R) and GABA. *Ex-vivo* slice recordings and *in-vivo* electrophysiological recordings from NCMs suggested that dopamine signals via D1R modulate the amplitude of GABAergic currents in NCM neurons and stimulus-specific neural plasticity. A similar modulation might occur in fly song preference learning, in which dopamine signals to pCd-2 *via* Dop1R2 possibly modulate GABAergic signals to pC1 neurons. It should be noted, however, that the present study does not clarify how each pCd-2 neuron expresses, and thus is modulated through, Rdl (GABA_A_) and Dop1R2 receptors, highlighting the importance of gene expression analysis at the single-cell resolution.

### Possible role of pCd-2 neurons in modulating the balance of mating-related decisions via pC1 neurons

When exposed to courtship songs, female flies typically escape from males before finally accepting copulation attempts (51). During this pre-copulation period, virgin females sometimes engage in aggressive behaviors (18). These aggressive behaviors towards the courting male are driven by pC1d/e neurons, while mating receptivity is promoted by other pC1 neurons (i.e., pC1a/b/c) to facilitate copulation acceptance (18). Our study shows that pC1d/e neurons receive synaptic inputs from intensive-type pCd-2 neurons (i.e., SMP286 and SMP287), while other pC1 neurons, pC1a/b/c, receive inputs from all pCd-2 neurons except SMP287 (Fig. 3 *G-J*). Because pCd-2 neurons are GABAergic, these pCd-2 neurons are likely to inhibit the target pC1 neurons, and thus adjust the balance between aggressive behaviors and copulation acceptance in female flies. Notably, two intensive-type pCd-2 neurons receive many synaptic inputs from pC1a/b/c neurons, suggesting that pC1a/b/c neurons suppress pC1d/e neurons via these pCd-2 neurons. Experiences of hearing conspecific songs might affect the landscape of pC1 activations, which would be mediated by mutual interactions between pCd-2 and pC1 neurons. In addition to reciprocal connections with pC1 neurons, pCd-2 neurons directly connect to the vpoDN (*SI Appendix*, Fig S6B). pCd-2 neurons thus modulate the activity landscape of the SAG-pC1-vpoDN pathway, which switches the female state from rejection to copulation acceptance: In this scenario, pCd-2 neurons are likely to play a role in inhibiting or gain-controlling this pathway if flies have prior auditory experiences. The EM database analysis also showed that one of the pCd-2 neurons, SMP286, receives direct synaptic inputs from circadian neurons involved in sleep regulation (i.e., LPN; see *SI Appendix*, Table S3) (52). These anatomical connections together imply that pCd-2 neurons serve as an integration hub for external sensory stimuli and internal states to achieve flexible control of mating receptivity.

### Perceptual learning in mammals, birds, and flies

‘Perceptual learning’ describes how perceptual discrimination ability improves with training (53, 54). A broad range of sensory skills improve with practice during perceptual learning, such as language acquisition, musical abilities, and visual discrimination (5, 55–58). Studies using animal models have shown that perceptual learning is associated with changes in sensory cortex activity (59–61). Notably, the involvement of GABA in perceptual learning is observed in the auditory cortex of Mongolian gerbils, in which the infusion of muscimol, a selective agonist for GABA_A_ receptors, prevents perceptual learning in the sound discrimination task (54). More peripherally, GABAergic granule cells in the olfactory bulb are responsible for olfactory perceptual learning of mice (62). These granule cells are the main source of lateral inhibition in the olfactory bulb and modulate pattern separation of mitral cells, which then project their axons to higher structures in the brain. Studies in this circuit further suggested the importance of inhibitory (i.e., granule cells) and excitatory (i.e., mitral cells) reciprocal connections in olfactory perceptual learning (62). The song preference learning observed in fruit flies also involves the inhibitory-excitatory reciprocal circuit that includes GABAergic signals through GABA_A_ receptors as a key motif, supporting the idea that the circuit mechanisms underlying this learning are shared between vertebrates and flies.

A “sensitive period”, which typically occurs during development including pre- and postnatal stages, has been identified during which the effect of experience on the brain to improve perceptual discrimination ability is particularly strong (63, 64). Studies using human and other vertebrate models have suggested that during the sensitive period, a balance between excitation and inhibition (E/I balance) is established, which typically requires appropriate sensory inputs (63–65). Particularly, the maturation of the inhibitory system plays a dominant role in shaping sensory perception during the sensitive period (66). In the primary auditory cortex of rodents, maturation of the inhibitory system during postnatal development enables excitation and inhibition to become highly correlated and balanced, which is accelerated by sound experiences (67). In the higher-level auditory cortex of songbird, GABAergic inhibitions matured by tutor song experiences form selective responses to the tutor song (11). Although the present study has not assessed the developmental aspect of song preference learning of flies, exploring the maturation process of the GABAergic signals to/from pCd-2 neurons in the fly model would provide key insights into a general neural-circuit mechanism on how sensory experiences establish the E/I balance that shapes information processing during development. An interesting future direction is to test if the neural circuit motif found in this study also underlies the experience-dependent auditory plasticity observed in vertebrates.

## Materials and Methods

### Fly stocks

*Drosophila melanogaster* were reared on standard yeast-based media at 25°C in 40% to 60% relative humidity and a 12-hour light/dark cycle. For fly strain details see Key Resources Table and *SI Appendix,* Table S1. The *dsx*∩*Gad1* driver was generated by chromosomal recombination of two hemi-drivers, *Gad1*^GAL4AD^ and *dsx*^GAL4DBD^. *Gad1*^GAL4AD^ replicates the expression pattern of *Glutamic acid decarboxylase 1* (*Gad1*), a gene encoding the rate-limiting enzyme in the GABA synthesis pathway, whereas *dsx*^GAL4DBD^ replicates the *dsx* expression pattern (24, 68). The *dsx*∩*Gad1*^brain^ driver was generated by combining the *dsx*∩*Gad1* driver with a flippase-mediated *GAL80* strain (69) which has two transgenes *Otd*-*FLP* (27) and *tubulin^P^ FRT-GAL80-FRT*. We used *Canton-S* as the wild-type strain. For knockdown experiments, *UAS-RNAi* strains, selected based on previous studies, were used with corresponding background strains as controls (from the Transgenic RNAi Project, see *SI Appendix* for details). All males used in the behavioral experiments were of the wild-type strain. They were collected within 8 h after eclosion, had their wings clipped under ice anesthesia, and kept singly until the copulation test was conducted as described previously (70).

### Sound stimulus

We used artificial pulse songs as described previously (70). The songs are comprised of repetitions of a 1-s pulse burst followed by a 2-s pause. Inter-pulse intervals are 35 ms for the conspecific song and 75 ms for the heterospecific song. Intrapulse frequency was 167 Hz in both songs. We delivered the sound stimuli using loudspeakers (FF225WK, FOSTEX, Foster Electric Company, Tokyo, Japan during training; Daito Voice AR-10N, Tokyo Cone Paper MFG. Co. Ltd in copulation tests). The mean baseline-to-peak amplitudes of the particle velocity were 6.6 to 8.6 mm/s during training and around 9.2 mm/s during the copulation test.

### Training

We define “training” as the process that exposes flies to external artificial sounds (22). Females were collected within 8 h after eclosion under ice anesthesia and kept in groups of 8 to 10 in training capsules at 24-26°C and 40-60% humidity. Training capsules are made from plastic straws about 70 mm long with a diameter of 14 mm, with edges covered with mesh and food provided at the bottom. In the experienced training condition, females were exposed to the conspecific song for 6 days. Naïve females were kept in training capsules for 6 days without song exposure. We replaced the capsule every 2-3 days (within 60 h) during the training. After training, females were kept without song exposure for 14-18 h and then subjected to the copulation test.

### Copulation test

We conducted the copulation test within 4 h after light onset (ZT 0 – 4) at 24-26°C and 40-60% humidity. 7-day-old wing-clipped males were used as a mating partner. A male and a female were gently aspirated into one of the eight chambers of the experimental plate (70) without anesthesia. The experimental plate was placed above a loudspeaker at a distance of 39 mm, and playback of sound stimuli started immediately. Fly pairs were recorded for 35 min at 15 fps with a web camera (Logicool HD Webcam C270, Tokyo, Japan). Copulation was defined as the following criteria: (1) the male mounts the female for more than 5 min, (2) the mounted female decreases locomotor activity, and (3) opens her wings during the mounting (15).

### Statistical analysis for behavioral analysis

Copulation test data were analyzed using R (version 4.1.0). We evaluated the cumulative copulation rates and copulation latencies (i.e., time to start copulation) of all groups simultaneously, using an accelerated failure time (AFT) model (25, 26) included in the flexsurv package (version 2.2.1) of R. The learning index (LI) and 95% confidence interval (CI) were estimated using log-logistic AFT model. Briefly, a lack of song preference learning results in a LI of 1, whereas LI > 1 indicates that flies demonstrate the learning phenotype (See *SI Appendix,* Supporting Information Text). To evaluate the cumulative copulation rate and copulation latency of each group, we also utilized restricted mean time lost (RMLT) (71) calculations using the survRM2 package (version 2.2.1) of R (15). Calculated p-values were adjusted by Bonferroni corrections.

### Immunohistochemical analysis

We performed immunohistochemistry as described previously (72). Briefly, the brains and ventral nerve cords of 5- to 7-day-old females were dissected in phosphate-buffered saline (PBS; pH 7.4), fixed for 1 h in 4% paraformaldehyde/PBS (TAKARA BIO INC. Cat#T900), and subjected to antibody labeling. Primary and secondary antibodies are listed in Key Resources Table.

### Confocal microscopy and image processing

Samples were imaged using an FV1200 laser-scanning confocal microscope (Olympus) equipped with either a silicone-oil immersion 30× or 60× objective lens (UPLSAPO 30×s, NA = 1.05; UPLSAPO 60×s, NA = 1.30; Olympus). Serial optical sections were obtained every 0.84 μm with a resolution of 512 × 512 pixels (0.83 μm /pixel). For three-dimensional (3D) image reconstruction, confocal image datasets were processed with VVDViewer (version 1.5.10). Image size, contrast, and brightness were adjusted using Fiji (version 2.9.0) (73).

### Detecting synaptic connections using the FlyEM database

GABAergic neurons that synapse onto pC1 neurons (*SI Appendix,* Table S2) were extracted by combining the Hemibrain v1.2.1 datasets (obtained in the FlyEM project; Scheffer et al., 2020) with a neurotransmitter classification tool (32). In order to identify pCd-2 neurons from the extracted GABAergic neuron population, we assessed whether cell bodies were located at the posterior-dorsal side of the brain and projected to the superior medial protocerebrum (SMP) and ipsilateral GNG, with a trajectory similar to that of pCd-2 neurons described in a previous study (19). Since the Hemibrain v1.2.1 dataset was derived from electron microscopy (EM) sections of a significant portion of the right hemibrain and a small portion of the left hemibrain, two pCd-2 neurons in the left hemibrain (i.e., SMP294 and SMP297) were not identified. The projection patterns of pCd-2 and pC1 neurons were visualized using Virtual Fly Brain (Figure 3*B*-*F*; (74). The neurons were registered to the JRC2018 template (75).

The number of synapses between two neurons was obtained from the NeuPrint web interface (https://neuprint.janelia.org/). In accordance with a previous study (18), paired neurons with 6 or more synapses connecting them were considered to be connected (*SI Appendix,* Table S3).

### Gene expression profile of pCd-2 neurons

To verify the gene expression profile of pCd-2 neurons, we analyzed a fly brain single-cell transcriptome dataset published previously (GEO accession: GSE107451; GSE107451_DGRP-551_w1118_WholeBrain_157k_0d_1d_3d_6d_9d_15d_30d_50d_10X_DGEM_MEX.mtx.tsv.tar.gz) (23) using Seurat package (ver. 4.3.0.1) for R (ver. 4.3.0). Expression levels were calculated using the NormalizeData function of Seurat with the LogNormalize method and default parameter settings (scale.factor = 10000). To characterize gene expression profiles of pCd-2 neurons, the 23 cells that express *elav* (a pan-neuronal marker), *Gad1*, and *dsx* were selected using the WhichCells function (parameter: expression > 0). The expression level of neurotransmitter receptor genes, including *rdl* for GABA_A_-type receptor-expressing cells and *dop1r1*, *dop1r2*, and *dopr2* as dopamine receptor-expressing cell markers of these neurons were analyzed.

## Acknowledgments

We thank Yusuke Miwa, Ken-ichi Kimura, and Daisuke Yamamoto for *Otd*-*FLP*; *tubulin^P^ FRT-GAL80-FRT* fly strain, Mako Murai, Takuro S Ohashi, Xiaodong Li, Yukina Chiba, Yoichi Oda, Matthew P Su, Naotoshi Nakamura, and Shingo Iwami for discussions, Bloomington Drosophila Stock Center for fly stocks, and Developmental Studies Hybridoma Bank for antibodies. This study was supported by MEXT KAKENHI Grants-in-Aid for Scientific Research (B) (Grant JP20H03355 to AK), Scientific Research on Innovative Areas “Evolinguistics” (Grant JP20H04997 to AK), Grant-in-Aid for Transformative Research Areas (A) “iPlasticity” (Grant JP21H05689 and JP23H04228 to AK), “Evolutionary theory for constrained and directional diversities” (Grant 20H04865 to YI), Grant-in-Aid for Early-Career Scientists (Grants JP21K15137 to RT), Grant-in-Aid for Transformative Research Areas (A) Hierarchical Bio-Navigation (Grant JP22H05650 to RT), JST PRESTO (JPMJPR21S2 to YI) and JST FOREST (Grant JPMJFR2147 to AK), Japan.

