## Supplemental Files for "Neural-circuit basis of song preference learning in fruit flies"

### Supporting Information Text

#### *dsx*∩*Gad1*$^{brain}$ neurons resemble pCd-2 neurons morphologically

*dsx*-expressing GABAergic neurons are found in the brain and ventral nerve chord (Figure 1A). Here we examined the morphology of the brain subset, which is labeled in *dsx*∩*Gad1*$^{brain}$ driver (Figure 2C). The labeled neurons in the brain, 4 neurons per hemibrain, have cell bodies ventral to the protocerebral bridge. From these cell bodies arise the neurites: The dorsal projection runs toward the anterior-medial part of the superior medial protocerebrum (SMP) with extensive arborizations before sending a long branch to the dorsal surface of the ipsilateral gnathal ganglia (GNG). The ventral projection runs anteroventrally and joins the aforementioned long branch to terminate in the ipsilateral GNG (Figure 2C). These morphological characteristics resemble those of pCd-2 neurons and are distinct from other *dsx*-expressing neuronal clusters in the female brain (1). Previous studies reported the number of pCd-2 neurons as 3 to 4 per hemibrain (1, 2), which also aligns with our observation of *dsx*-expressing GABAergic neurons in the brain.

#### The genetic background of fly strains affects song responses

In wild-type flies (*Canton-S* strain), naïve females, who have never been exposed to courtship songs, showed strong behavioral responses to both the conspecific song and the heterospecific song (*SI Appendix*, Fig. S1). This phenotype was typically observed in most of the control groups that were used as the background strain for RNAi knockdown (Figures 1, 2). However, naïve females in some control groups showed a reduced response to the heterospecific song as compared to the conspecific song (Fig. 4*C* left). We therefore introduce the AFT model in this study to assess the behavioral phenotype: If the difference in the responses to the two songs becomes larger after song experience, we regarded the fly group as showing song preference learning. To this end, we evaluated the interaction of song experience and test song type using the AFT model (see the following section).

#### Data analysis using the AFT model

We evaluated the cumulative copulation rates and copulation latencies of all of the four groups (i.e., NC, NH, EC, and EH in Figures 1, 2, and 4) simultaneously, using an accelerated failure time (AFT) model (3, 4). Prior to analysis, we tested several distributions to formulate the AFT model framework with our datasets. We tested Weibull, exponential, log-normal, normal, logistic, and log-logistic distributions and found that the log-logistic distribution gave the smallest values for both the Akaike's Information Criterion (AIC) and Bayesian Information Criterion (BIC) in all datasets. Therefore, the log-logistic distribution was used throughout the analyses.

The AFT model assesses the effect of covariates or interactions on the time to copulate (i.e., copulation latency), $T$. In this study, we defined two types of covariates: $\chi_1$, the existence of conspecific song exposure in the training session (naïve *vs.* experienced), and $\chi_2$, the test song type (conspecific song *vs.* heterospecific song). Since each covariate has two factors, we set one factor as *0* and the other as *1* for each covariate. The effect of the interaction between covariates $\chi_1$ and $\chi_2$ is considered to be the effect of song preference learning. Using the log-logistic AFT model, copulation latency $T$ is described as follows:

$$T = e^{(\beta_0+\beta_1\chi_1+\beta_2\chi_2+\gamma\chi_1\chi_2+\sigma\varepsilon)}$$

where $\varepsilon$ is the residual that corresponds to log-logistic distribution and $\sigma$ is the scale parameter. To assess song preference learning, we focused on the interaction of the two covariates, $\chi_1$ and $\chi_2$. In particular, $e^{\gamma}$, an acceleration factor of the interaction, was used as the learning Index, LI. The results of the analysis are described in Table Supplemental 4.

**Fig. S1**. **Song preference learning in wild-type flies**

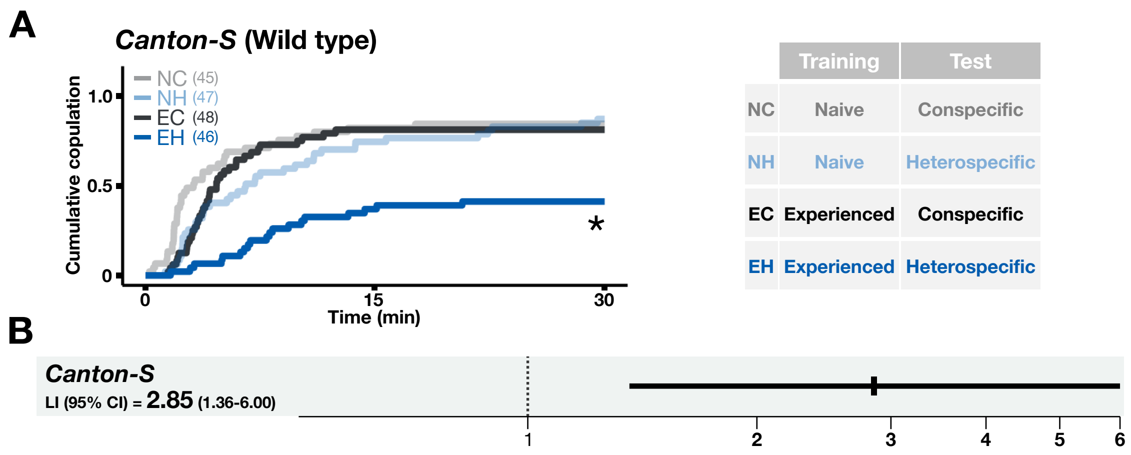


(A) Cumulative copulation rates of wild-type flies (*Canton-S* strain). NC, naïve flies tested with the conspecific song; NH, naïve flies tested with heterospecific song.; EC, experienced flies tested with the conspecific song; EH, experienced flies tested with heterospecific song. The number of trials in each group is shown in parentheses. (B) Learning index (LI) of wild-type flies. Square and line indicate estimated LI and 95% confidence interval (CI), respectively. *p<0.05; the log-logistic AFT model.

**Fig. S2. *Gad1* knockdown with *Gad1-TRiP-RNAi attP2* strain**

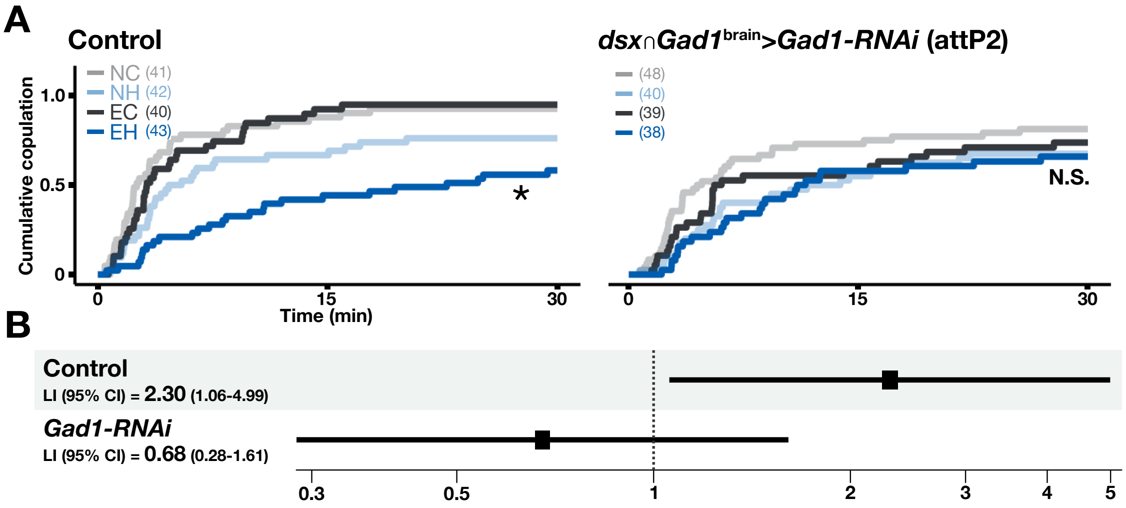


(A) Cumulative copulation rates in control and pCd-2 specific *Gad1* knockdown groups in *Gad1-TRiP-RNAi attP2* strain (See Table supplement 1 for genotypes). The *dsx*∩*Gad1*$^{brain}$ driver was used. NC, naïve flies tested with the conspecific song; NH, naïve flies tested with heterospecific song.; EC, experienced flies tested with the conspecific song; EH, experienced flies tested with heterospecific song. The number of trials in each group is shown in parentheses. N.S., not significant; *p<0.05; Log-logistic AFT model. (B) Learning index (LI) with a 95% confidence interval (CI).

**Fig. S3. GABA knockdown by the *dsx*∩*Gad1*$^{brain2}$ driver disturbs song preference learning**

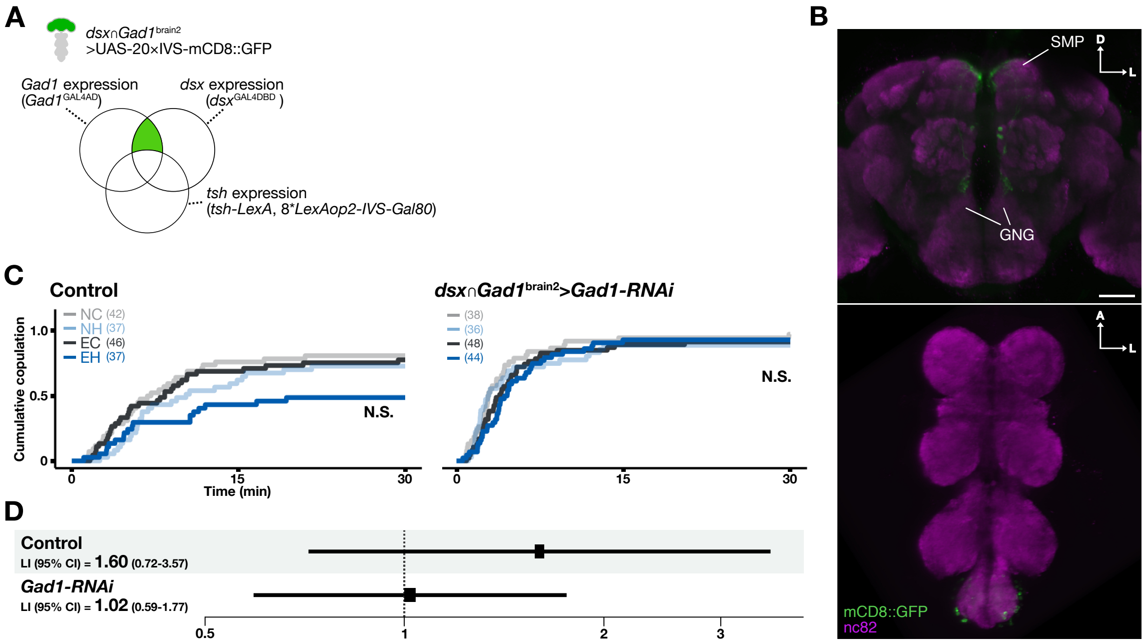


(A) Venn diagram of genetic intersection. The area containing *dsx*∩*Gad1*$^{brain2}$ -labeled neurons is marked in green. (B) The *dsx*∩*Gad1*$^{brain2}$ expression in females. The central brain (top; anterior view) and ventral nerve chord (bottom; ventral view) are shown. Scale bars, 50 µm. A, anterior; D, dorsal; L, Lateral (the same in the following figures). (C) Cumulative copulation rates in control and *dsx*∩*Gad1*$^{brain2}$ > *Gad1* knockdown groups. NC, naïve flies tested with the conspecific song; NH, naïve flies tested with heterospecific song; EC, experienced flies tested with the conspecific song; EH, experienced flies tested with heterospecific song. The number of trials in each group is shown in parentheses. N.S., not significant; *p<0.05; Log-logistic AFT model. (D) Learning index (LI) with a 95% confidence interval (CI).

#### Fig. S4. Synaptic sites of pCd-2 neurons

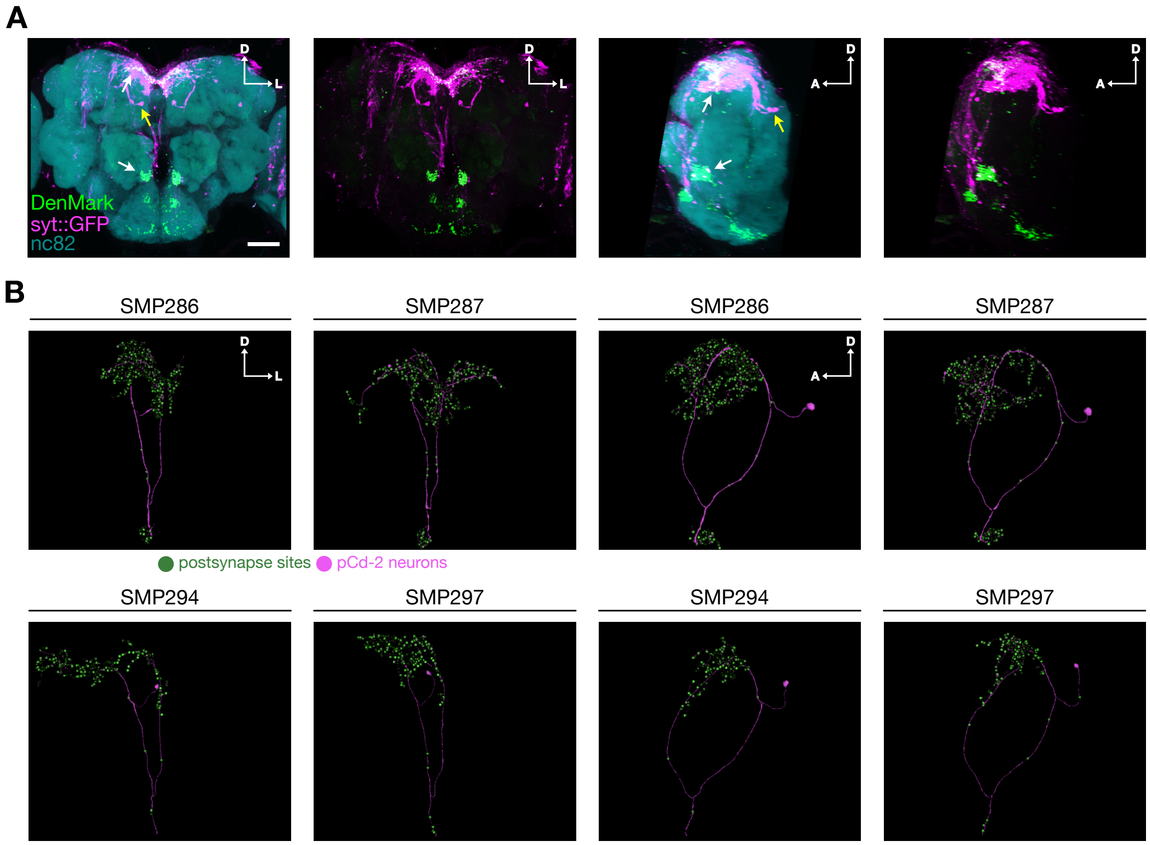


(A) Distribution of the postsynaptic marker in pCd-2 neurons. The *dsx*∩*Gad1* driver was used to express DenMark (green) and syt::GFP (magenta), which are supposed to localize at presynaptic and postsynaptic sites respectively. Although DenMark signals were localized only in specific regions in the brain (white arrows), syt::GFP signals were distributed throughout pCd-2 neurons including cell bodies (yellow arrow) without specific localized regions possibly due to excess expression. Scale bar, 50 $\mu$m. (B) Distribution of the postsynaptic sites on four pCd-2 neurons in the FlyEM dataset. Image was obtained from NeuPrint (https://neuprint.janelia.org/) based on the Hemibrain v1.2.1 dataset. Frontal views (left two panels) and lateral views (right two panels) are shown.

**Fig. S5. pC1 neurons that synapse with each pCd-2 neuron**

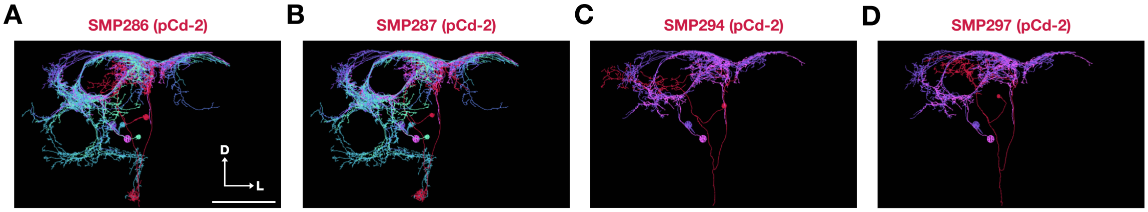


pCd-2 neuron (Red) and its connecting pC1 neurons (magenta, purple, blue, light blue, and spring green). SMP286 (A), SMP287 (B), SMP294 (C), and SMP297 (D) in the right hemibrain are overlaid with pC1 neurons using Virtual Fly Brain (https://www.virtualflybrain.org). The color of each pC1 neuron corresponds to that used in Figure 3 *F-I*. Scale bar, 50µm.

**Fig. S6. Synaptic connections between pCd-2 neurons, neurons involved in female copulation receptivity, and dopaminergic neurons**

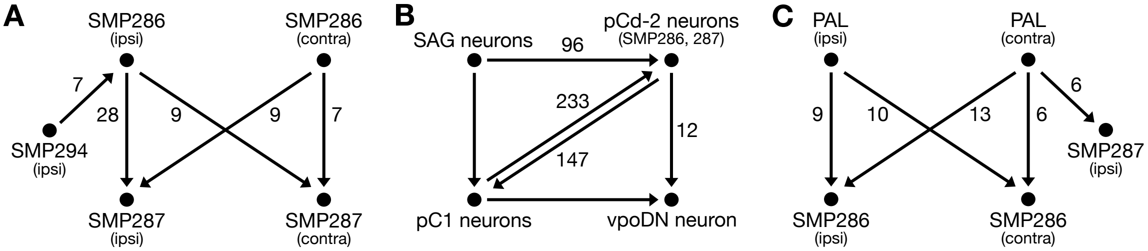


(A) Synaptic connections within pCd-2 neurons (indicated by arrows). Numbers on arrows indicate the number of synapses. (B) Synaptic connections with neurons involved in copulation receptivity. Only the right-hemibrain data are shown. The right hemibrain contains two SAG neurons, five pC1 neurons, and one vpoDN neuron in the Hemibrain v1.2.1 dataset (https://neuprint.janelia.org/). The total synapse number between two pCd-2 neurons (SMP286 and SMP287) and SAG neurons, pC1 neurons, or vpoDN neuron located in the right hemibrain is indicated. (C) Synaptic connections with PAL dopaminergic neurons. ipsi, ipsilateral; contra, contralateral. To count the number of synapses, we only took into account synaptic connections with 6 or more synapses between paired neurons (A-C).

**Fig. S7. Expression levels of *Rdl* and *dopamine receptor* genes in dsx+/gad1+/elav+ cells**

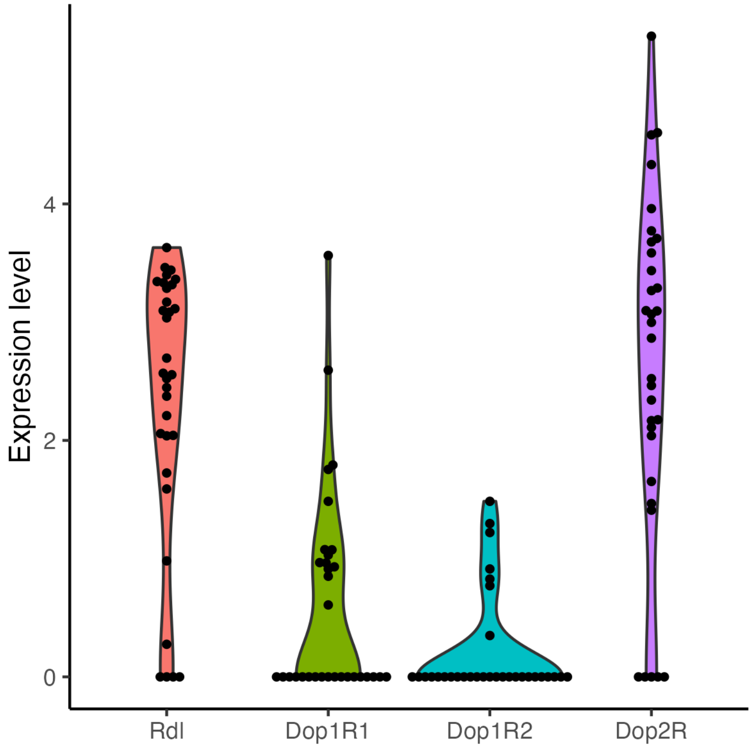


Expression levels of each receptor (*Rdl, dop1r1*, *dop1r2*, and *dopr2*) gene in pCd-2 neurons from a single-cell transcriptome data (See Materials and Methods). Each dot indicates the expression level in a single dsx+/gad1+/elav+ cell (N = 23 cells). This single-cell RNA sequence dataset was obtained from 40 flies (*D. melanogaster* adult brains, 20 females and 20 males)(5).

**Fig. S8. Knockdown of two dopamine receptors**

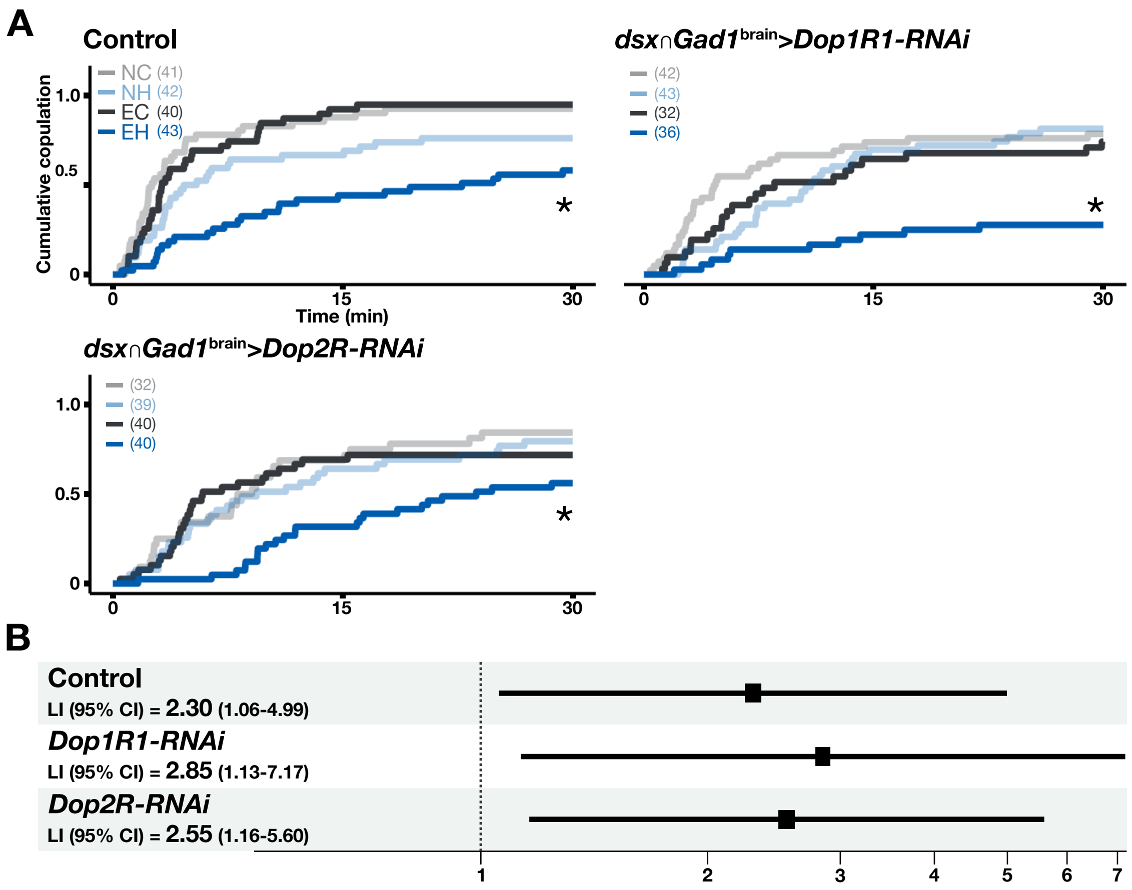


(A) Cumulative copulation rates in control and dopamine receptor (*Dop1R1* or *Dop2R*) knockdown groups. The *dsx*∩*Gad1*$^{\text{brain}}$ driver was used. NC, naïve flies tested with the conspecific song; NH, naïve flies tested with heterospecific song; EC, experienced flies tested with the conspecific song; EH, experienced flies tested with heterospecific song.*p<0.05; Log-logistic AFT model. (B) Learning index (LI) with a 95% confidence interval (CI) in the control and knockdown groups.

**Fig. S9. Labeling pattern of the *dsx*∩*Gad1* driver in the male brain**

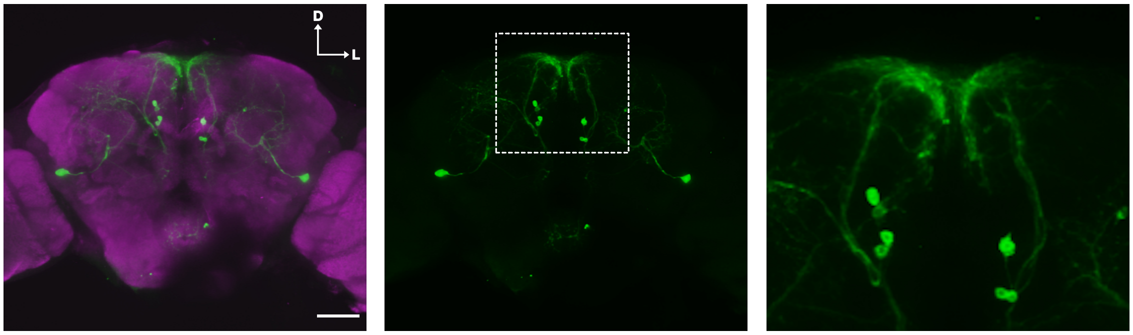


Overview of the central brain (left and middle; posterior view) and a magnified view (right) of the neurites and cell bodies located in the superior medial protocerebrum (SMP) are shown. Expression was visualized with GFP (green). Brain neuropils were labeled with nc82 (magenta). Scale bars, 50 µm.

#### Table S1. Fly genotypes

| Figure | genotype |
|---|---|
| 1A | w+;+;20xUAS-IVS-mCD8::GFP(attP2)/Gad1$^{GAL4AD}$, dsx$^{GAL4DBD}$ |
| 1D,E | w+;CaryP(attp40);Gad1$^{GAL4AD}$,dsx$^{GAL4DBD}$ |
| | w+;UAS-Gad1-RNAi(attp40);Gad1$^{GAL4AD}$, dsx$^{GAL4DBD}$ |
| 2B,C | w+;Otd-FLP,tubP-FRP-GAL80-FRT;20xUAS-IVS-mCD8::GFP(attP2)/Gad1$^{GAL4AD}$,dsx$^{GAL4DBD}$ |
| 2D,E | w+;Otd-FLP,tubP-FRP-GAL80-FRT/CaryP(attp40);Gad1$^{GAL4AD}$,dsx$^{GAL4DBD}$ |
| | w+;Otd-nls:FLPo,tubP-FRP-GAL80-FRT/UAS-Gad1-RNAi(attp40);Gad1$^{GAL4AD}$,dsx$^{GAL4DBD}$ |
| 3A | w[1118];R71G01-lexA(attP40)/lexAopCD4::spGFP11;GFP1~10,GFP11; Gad1$^{GAL4AD}$,dsx$^{GAL4DBD}$/UAS-CD4::spGFP1-10 |
| 4A,B | w+;Otd-nls:FLPo,tubP-FRP-GAL80-FRT/CaryP(attP40);Gad1$^{GAL4AD}$,dsx$^{GAL4DBD}$ |
| | w+; Otd-nls:FLPo,tubP-FRP-GAL80-FRT/UAS-Rdl-RNAi(attP40);Gad1$^{GAL4AD}$,dsx$^{GAL4DBD}$ |
| 4C,D | w+;Otd-nls:FLPo,tubP-FRP-GAL80-FRT;CaryP(attP2)/Gad1$^{GAL4AD}$,dsx$^{GAL4DBD}$ |
| | w+;Otd-nls:FLPo,tubP-FRP-GAL80-FRT; UAS-Dop1R2-RNAi(attP2)/Gad1$^{GAL4AD}$,dsx$^{GAL4DB}$ |
| S1 | Canton-S |
| S2 | w+;Otd-FLP,tubP-FRP-GAL80-FRT;CaryP(attp2)/Gad1$^{GAL4AD}$,dsx$^{GAL4DBD}$ |
| | w+;Otd-nls:FLPo,tubP-FRP-GAL80-FRT;UAS-Gad1-RNAi(attp2)/Gad1$^{GAL4AD}$,dsx$^{GAL4DBD}$ |
| S3 | w+;tsh-LexA, pJFRC20-8xLexAop2-IVS-GAL80-WPRE(su(Hw)attp5)/CaryP(attp40); Gad1$^{GAL4AD}$,dsx$^{GAL4DBD}$ |
| | w+;tsh-LexA, pJFRC20-8xLexAop2-IVS-GAL80-WPRE(su(Hw)attp5)/UAS-Gad1-RNAi(attp40); Gad1$^{GAL4AD}$,dsx$^{GAL4DBD}$ |
| S4A | w+;UAS-DenMark, UAS-syt.eGFP;Gad1$^{GAL4AD}$,dsx$^{GAL4DB}$ |
| S8A,B | w+;Otd-nls:FLPo,tubP-FRP-GAL80-FRT;CaryP(attP2)/Gad1$^{GAL4AD}$,dsx$^{GAL4DBD}$ |
| | w+;Otd-nls:FLPo,tubP-FRP-GAL80-FRT; UAS-Dop1R1-RNAi(attP2)/Gad1$^{GAL4AD}$,dsx$^{GAL4DB}$ |
| | w+;Otd-nls:FLPo,tubP-FRP-GAL80-FRT; UAS-Dop2R-RNAi(attP2)/Gad1$^{GAL4AD}$,dsx$^{GAL4DB}$ |
| S9 | w+;+;20xUAS-IVS-mCD8::GFP(attP2)/Gad1$^{GAL4AD}$, dsx$^{GAL4DBD}$ |

#### **Table S2.** List of neurons that have GABA synapses to pC1 neurons

| Neuron ID | body name |
|---|---|
| 611413241 | (ADM04)_L |
| 328377109 | (aDT6e)(MBDLaxon2) |
| 5813047422 | (aDT6e)(MBDLaxon2) |
| 328377109 | (aDT6e)(MBDLaxon2) |
| 548972278 | (aDT6f)(MBDLaxon2) |
| 642422617 | (MBDLaxon1) |
| 828282250 | (MBDLaxon1) |
| 487286529 | (MBDLaxon1) |
| 1137862716 | (pIP5d) |
| 1038070642 | AOTU064(SCB012)_R |
| 1629383757 | Ascending? |
| 1197449936 | aSP8(aSP8a)_R |
| 1166416004 | AVLP008_R |
| 1013046019 | AVLP008_R |
| 1044075948 | AVLP008_R |
| 580227374 | AVLP029_R |
| 1166545210 | AVLP095_R |
| 1107441034 | AVLP096_R |
| 1200554295 | AVLP096_R |
| 1322051647 | AVLP255_R |
| 922595498 | AVLP256_R |
| 1292971921 | AVLP256_R |
| 5813047489 | AVLP531_R |
| 800579317 | CL037_R |
| 516055524 | CL063_R |
| 510602264 | CL125_R |
| 5813021221 | CL125_R |
| 5813040037 | CL176_R |
| 666252297 | CL234_R |
| 5812980507 | CL234_R |
| 1788237005 | LAL059_R |
| 5901226003 | LAL059_R |
| 485934965 | oviIN_L |
| 423101189 | oviIN_R |
| 5812990346 | pC2l(PVL06)_L |
| 1501392572 | PS088_R |
| 1352685625 | PVLP047_R |
| 1568596284 | PVLP047_R |
| 1598586139 | PVLP047_R |
| 1598944549 | PVLP047_R |
| 1414366899 | PVLP093(PDL09)_L |
| 1538562093 | PVLP130(PVL06)_L |
| 517587356 | SAG |
| 5812981862 | SAG |
| 581409749 | SMP093(ADM04)_L |
| 425143700 | SMP097_a_R |

| Neuron ID | body name |
|---|---|
| 733458828 | SMP163_R |
| 611693656 | SMP164_R |
| 328373131 | SMP286(PDM05)_L |
| 297947227 | SMP286_R |
| 5813068453 | SMP287(PDM05)_L |
| 5813069072 | SMP287_R |
| 389563849 | SMP294_R |
| 5813071288 | SMP297_R |
| 1409129636 | VES020(AVM11)_L |
| 1136852717 | WED014_R |
| 2067847820 | WED014_R |
| 390774986 | |
| 421137065 | |
| 422151685 | |
| 297925628 | |
| 451485030 | |
| 639694375 | |
| 360081309 | |
| 946619892 | |
| 855673094 | |
| 983025411 | |
| 1136852435 | |
| 1321710747 | |
| 1353423473 | |
| 1507592058 | |
| 1600334745 | |
| 1662092808 | |
| 946619892 | |
| 704816317 | |
| 5813083315 | |

#### Table S3. List of neurons that output to or input from pCd-2 neurons

| Neron ID (Output to pCd-2) | Neuron name | Corresponding pCd-2 subtype | Synapse number | Neuron ID (Input from pCd-2) | Neuron name | Corresponding pCd-2 subtype | Synapse number |
|---|---|---|---|---|---|---|---|
| 5813046951 | pC1a | SMP286(ID: 297947227) | 119 | 297580589 | SMP548 | SMP286_R(297947227) | 42 |
| 386833850 | SMP529 | SMP286(297947227) | 74 | 5813046951 | pC1a | SMP286_R(297947227) | 38 |
| 327937506 | | SMP286(297947227) | 54 | 267214250 | pC1b | SMP286_R(297947227) | 32 |
| 388638672 | SMP161 | SMP286(297947227) | 45 | 5813063587 | pC1d | SMP286_R(297947227) | 28 |
| 453527730 | SMP161 | SMP286(297947227) | 43 | 5813069072 | SMP287 | SMP286_R(297947227) | 28 |
| 419345606 | SMP202 | SMP286(297947227) | 41 | 5812980529 | SLP212 | SMP286_R(297947227) | 24 |
| 298262408 | | SMP286(297947227) | 40 | 392821837 | pC1b | SMP286_R(297947227) | 21 |
| 357949102 | SMP222 | SMP286(297947227) | 37 | 452689494 | SMP550 | SMP286_R(297947227) | 20 |
| 297925887 | | SMP286(297947227) | 37 | 267551639 | pC1c | SMP286_R(297947227) | 17 |
| 297921608 | SMP333 | SMP286(297947227) | 37 | 328611467 | SMP480 | SMP286_R(297947227) | 16 |
| 359744514 | pC1a | SMP286(297947227) | 35 | 326901620 | SMP288 | SMP286_R(297947227) | 15 |
| 297584869 | SMP035 | SMP286(297947227) | 34 | 297921608 | SMP333 | SMP286_R(297947227) | 15 |
| 327596481 | SMP035 | SMP286(297947227) | 32 | 2225880115 | | SMP286_R(297947227) | 14 |
| 542747457 | SLP324 | SMP286(297947227) | 27 | 5813009651 | SMP480 | SMP286_R(297947227) | 13 |
| 357940977 | SMP218 | SMP286(297947227) | 27 | 5813055769 | SLP212 | SMP286_R(297947227) | 13 |
| 328611467 | SMP480 | SMP286(297947227) | 27 | 2132434539 | | SMP286_R(297947227) | 12 |
| 328632707 | SMP083 | SMP286(297947227) | 26 | 359744514 | pC1a | SMP286_R(297947227) | 12 |
| 297930119 | | SMP286(297947227) | 24 | 5813057864 | vpoDN | SMP286_R(297947227) | 12 |
| 425143700 | SMP097_a | SMP286(297947227) | 23 | 297929815 | SMP480 | SMP286_R(297947227) | 12 |
| 417204164 | SLP324 | SMP286(297947227) | 23 | 5813011471 | SMP479 | SMP286_R(297947227) | 12 |
| 358281935 | SMP218 | SMP286(297947227) | 23 | 390331583 | SMP368 | SMP286_R(297947227) | 12 |
| 5813009845 | SMP218 | SMP286(297947227) | 22 | 2132775255 | | SMP286_R(297947227) | 11 |
| 543566744 | SMP540 | SMP286(297947227) | 22 | 5813008911 | SMP486_d | SMP286_R(297947227) | 11 |
| 390702220 | | SMP286(297947227) | 22 | 327937506 | | SMP286_R(297947227) | 11 |
| 357923095 | SMP539 | SMP286(297947227) | 22 | 2255192411 | | SMP286_R(297947227) | 10 |
| 448515267 | SMP221 | SMP286(297947227) | 20 | 298961895 | SMP478 | SMP286_R(297947227) | 10 |
| 358290805 | SMP083 | SMP286(297947227) | 20 | 358635284 | SMP480 | SMP286_R(297947227) | 10 |
| 328010530 | SMP454 | SMP286(297947227) | 20 | 5813026589 | SMP082 | SMP286_R(297947227) | 9 |
| 5813010435 | SMP540 | SMP286(297947227) | 19 | 2226221475 | | SMP286_R(297947227) | 9 |
| 486932201 | SMP083 | SMP286(297947227) | 19 | 5813068453 | SMP287 | SMP286_R(297947227) | 9 |
| 297929815 | SMP480 | SMP286(297947227) | 19 | 5813003676 | | SMP286_R(297947227) | 8 |
| 297588882 | | SMP286(297947227) | 19 | 2162779161 | | SMP286_R(297947227) | 8 |
| 5813069630 | NPFL1-I | SMP286(297947227) | 18 | 328265389 | SMP082 | SMP286_R(297947227) | 8 |
| 976934253 | SMP594 | SMP286(297947227) | 18 | 2101399263 | | SMP286_R(297947227) | 8 |
| 581690438 | SMP097_a | SMP286(297947227) | 18 | 423101189 | oviIN | SMP286_R(297947227) | 8 |

| | | | | | | | |
|---|---|---|---|---|---|---|---|
| 297925742 | SMP480 | SMP286(297947227) | 18 | 514850616 | pC1e | SMP286_R(297947227) | 8 |
| 545906392 | | SMP286(297947227) | 17 | 5813026613 | SMP530 | SMP286_R(297947227) | 8 |
| 514842241 | SMP253 | SMP286(297947227) | 16 | 297921642 | SMP486_b | SMP286_R(297947227) | 8 |
| 480521586 | | SMP286(297947227) | 16 | 425143700 | SMP097_a | SMP286_R(297947227) | 7 |
| 327588225 | SMP083 | SMP286(297947227) | 16 | 423442357 | SMP480 | SMP286_R(297947227) | 7 |
| 298266950 | | SMP286(297947227) | 16 | 545906392 | | SMP286_R(297947227) | 7 |
| 296859438 | SMP297 | SMP286(297947227) | 16 | 5813092319 | SMP346 | SMP286_R(297947227) | 7 |
| 540329317 | SLP324 | SMP286(297947227) | 15 | 514842241 | SMP253 | SMP286_R(297947227) | 7 |
| 513848395 | | SMP286(297947227) | 15 | 360254108 | SMP334 | SMP286_R(297947227) | 7 |
| 326253554 | SMP454 | SMP286(297947227) | 15 | 519694628 | SMP333 | SMP286_R(297947227) | 7 |
| 297921642 | SMP486_b | SMP286(297947227) | 15 | 297925742 | SMP480 | SMP286_R(297947227) | 7 |
| 5813008892 | | SMP286(297947227) | 14 | 5813055766 | SMP486_b | SMP286_R(297947227) | 7 |
| 5812981862 | SAG | SMP286(297947227) | 14 | 423792217 | SMP121 | SMP286_R(297947227) | 7 |
| 518930220 | SMP097_b | SMP286(297947227) | 14 | 390702220 | | SMP286_R(297947227) | 7 |
| 514527219 | | SMP286(297947227) | 14 | 298288301 | SMP481 | SMP286_R(297947227) | 6 |
| 419665356 | | SMP286(297947227) | 14 | 487623053 | | SMP286_R(297947227) | 6 |
| 299954300 | SIP076 | SMP286(297947227) | 14 | 2223144438 | | SMP286_R(297947227) | 6 |
| 5813055817 | PAL01 | SMP286(297947227) | 13 | 636944318 | SMP383 | SMP286_R(297947227) | 6 |
| 5813009950 | | SMP286(297947227) | 13 | 298262408 | | SMP286_R(297947227) | 6 |
| 5813009589 | | SMP286(297947227) | 13 | 298266797 | | SMP286_R(297947227) | 6 |
| 517587356 | SAG | SMP286(297947227) | 13 | 327933208 | SMP486_f | SMP286_R(297947227) | 6 |
| 480029788 | LPN | SMP286(297947227) | 13 | 327255181 | SMP290 | SMP286_R(297947227) | 6 |
| 450034902 | LPN | SMP286(297947227) | 13 | 297588775 | SMP481 | SMP286_R(297947227) | 6 |
| 423792440 | SMP097_b | SMP286(297947227) | 13 | 297588838 | SMP486_b | SMP286_R(297947227) | 6 |
| 387503511 | SLP324 | SMP286(297947227) | 13 | 359744514 | pC1a | SMP286(328373131) | 33 |
| 298258611 | SLP388 | SMP286(297947227) | 13 | 393499533 | SMP548 | SMP286(328373131) | 28 |
| 703145276 | | SMP286(297947227) | 12 | 392821837 | pC1b | SMP286(328373131) | 22 |
| 581055513 | | SMP286(297947227) | 12 | 5813055769 | SLP212 | SMP286(328373131) | 20 |
| 420010560 | | SMP286(297947227) | 12 | 267214250 | pC1b | SMP286(328373131) | 20 |
| 389568268 | SLP139 | SMP286(297947227) | 12 | 545621587 | | SMP286(328373131) | 17 |
| 5813055766 | SMP486_b | SMP286(297947227) | 11 | 519694628 | SMP333 | SMP286(328373131) | 17 |
| 5813019568 | | SMP286(297947227) | 11 | 487623053 | | SMP286(328373131) | 14 |
| 2317954019 | | SMP286(297947227) | 11 | 5812980529 | SLP212 | SMP286(328373131) | 14 |
| 633447561 | AVLP472 | SMP286(297947227) | 11 | 423852404 | | SMP286(328373131) | 13 |
| 580348303 | | SMP286(297947227) | 11 | 485934965 | oviIN | SMP286(328373131) | 13 |
| 604889732 | | SMP286(297947227) | 11 | 5813046951 | pC1a | SMP286(328373131) | 12 |
| 580719155 | | SMP286(297947227) | 11 | 297925742 | SMP480 | SMP286(328373131) | 12 |
| 422837964 | | SMP286(297947227) | 11 | 298961895 | SMP478 | SMP286(328373131) | 12 |
| 359326692 | SMP509 | SMP286(297947227) | 11 | 581405615 | | SMP286(328373131) | 12 |

| | | | | | | | |
|---|---|---|---|---|---|---|---|
| 580032546 | | SMP286(297947227) | 10 | 703145276 | | SMP286(328373131) | 11 |
| 520272258 | SMP098_a | SMP286(297947227) | 10 | 391125788 | | SMP286(328373131) | 11 |
| 516944116 | | SMP286(297947227) | 10 | 328709642 | | SMP286(328373131) | 10 |
| 420347328 | | SMP286(297947227) | 10 | 577330676 | SMP383 | SMP286(328373131) | 10 |
| 417143726 | SMP223 | SMP286(297947227) | 10 | 487636204 | SMP478 | SMP286(328373131) | 10 |
| 357944930 | SMP539 | SMP286(297947227) | 10 | 452849628 | | SMP286(328373131) | 9 |
| 297921808 | SIP076 | SMP286(297947227) | 10 | 5813069072 | SMP287 | SMP286(328373131) | 9 |
| 666567261 | AVLP472 | SMP286(297947227) | 9 | 550319575 | pC1c | SMP286(328373131) | 9 |
| 581055643 | | SMP286(297947227) | 9 | 486604771 | | SMP286(328373131) | 8 |
| 571692839 | SLP324 | SMP286(297947227) | 9 | 5813108370 | | SMP286(328373131) | 8 |
| 518934633 | | SMP286(297947227) | 9 | 267551639 | pC1c | SMP286(328373131) | 8 |
| 519694628 | SMP333 | SMP286(297947227) | 9 | 391121098 | | SMP286(328373131) | 7 |
| 485511772 | PAL01 | SMP286(297947227) | 9 | 5813068453 | SMP287 | SMP286(328373131) | 7 |
| 455172836 | SMP482 | SMP286(297947227) | 9 | 424879778 | | SMP286(328373131) | 6 |
| 358286442 | SMP466 | SMP286(297947227) | 9 | 391124880 | | SMP286(328373131) | 6 |
| 361312146 | SMP103 | SMP286(297947227) | 9 | 298262408 | | SMP286(328373131) | 6 |
| 387170738 | SLP324 | SMP286(297947227) | 9 | 424193645 | | SMP286(328373131) | 6 |
| 298586567 | | SMP286(297947227) | 9 | 514527219 | | SMP286(328373131) | 6 |
| 297925628 | | SMP286(297947227) | 9 | 5901195945 | SMP121 | SMP286(328373131) | 6 |
| 298599269 | | SMP286(297947227) | 9 | 517578770 | SMP098_a | SMP286(328373131) | 6 |
| 5813010533 | | SMP286(297947227) | 8 | 360768754 | | SMP286(328373131) | 6 |
| 5813009651 | SMP480 | SMP286(297947227) | 8 | 329310230 | SMP082 | SMP286(328373131) | 6 |
| 762128241 | | SMP286(297947227) | 8 | 486544396 | PAL02 | SMP286(328373131) | 6 |
| 612777140 | | SMP286(297947227) | 8 | 361351171 | SMP028 | SMP287(5813069072) | 19 |
| 580369597 | | SMP286(297947227) | 8 | 5813063587 | pC1d | SMP287(5813069072) | 16 |
| 579014562 | SMP103 | SMP286(297947227) | 8 | 330661529 | SMP172 | SMP287(5813069072) | 14 |
| 573151229 | | SMP286(297947227) | 8 | 2224162442 | | SMP287(5813069072) | 13 |
| 578642656 | | SMP286(297947227) | 8 | 581405615 | | SMP287(5813069072) | 12 |
| 571709547 | SMP233 | SMP286(297947227) | 8 | 362395402 | SMP172 | SMP287(5813069072) | 10 |
| 550388353 | | SMP286(297947227) | 8 | 298961895 | SMP478 | SMP287(5813069072) | 9 |
| 516943720 | | SMP286(297947227) | 8 | 704487840 | SMP602 | SMP287(5813069072) | 8 |
| 486255488 | | SMP286(297947227) | 8 | 488231939 | SMP093 | SMP287(5813069072) | 8 |
| 455513316 | | SMP286(297947227) | 8 | 5813020993 | SIP067 | SMP287(5813069072) | 8 |
| 361351171 | SMP028 | SMP286(297947227) | 8 | 487636204 | SMP478 | SMP287(5813069072) | 8 |
| 329732855 | SMP368 | SMP286(297947227) | 8 | 5813025545 | PI1_b | SMP287(5813069072) | 7 |
| 327588074 | | SMP286(297947227) | 8 | 393171256 | SMP172 | SMP287(5813069072) | 7 |
| 327242169 | SMP035 | SMP286(297947227) | 8 | 2162779159 | | SMP287(5813069072) | 7 |
| 298275090 | | SMP286(297947227) | 8 | 297925719 | SMP478 | SMP287(5813069072) | 7 |
| 5813060687 | | SMP286(297947227) | 7 | 2162779161 | | SMP287(5813069072) | 6 |

| | | | | | | | |
|---|---|---|---|---|---|---|---|
| 5813055759 | SMP467 | SMP286(297947227) | 7 | 327255181 | SMP290 | SMP287(5813069072) | 6 |
| 5813013636 | | SMP286(297947227) | 7 | 2194845024 | | SMP287(5813069072) | 6 |
| 5813011858 | | SMP286(297947227) | 7 | 5812980467 | | SMP287(5813069072) | 6 |
| 5813010120 | SMP301 | SMP286(297947227) | 7 | 2226221475 | | SMP287(5813069072) | 6 |
| 2286568672 | | SMP286(297947227) | 7 | 361351171 | SMP028 | SMP287(5813068453) | 19 |
| 1141035086 | | SMP286(297947227) | 7 | 5813063587 | pC1d | SMP287(5813068453) | 16 |
| 697645622 | SMP467 | SMP286(297947227) | 7 | 330661529 | SMP172 | SMP287(5813068453) | 14 |
| 610381395 | | SMP286(297947227) | 7 | 2224162442 | | SMP287(5813068453) | 13 |
| 548660840 | | SMP286(297947227) | 7 | 581405615 | | SMP287(5813068453) | 12 |
| 518593663 | | SMP286(297947227) | 7 | 362395402 | SMP172 | SMP287(5813068453) | 10 |
| 519017242 | | SMP286(297947227) | 7 | 298961895 | SMP478 | SMP287(5813068453) | 9 |
| 483880436 | | SMP286(297947227) | 7 | 704487840 | SMP602 | SMP287(5813068453) | 8 |
| 451485030 | | SMP286(297947227) | 7 | 488231939 | SMP093 | SMP287(5813068453) | 8 |
| 418641700 | SMP537 | SMP286(297947227) | 7 | 5813020993 | SIP067 | SMP287(5813068453) | 8 |
| 361105354 | | SMP286(297947227) | 7 | 487636204 | SMP478 | SMP287(5813068453) | 8 |
| 390331583 | SMP368 | SMP286(297947227) | 7 | 5813025545 | PI1_b | SMP287(5813068453) | 7 |
| 389563849 | SMP294 | SMP286(297947227) | 7 | 393171256 | SMP172 | SMP287(5813068453) | 7 |
| 329728593 | | SMP286(297947227) | 7 | 2162779159 | | SMP287(5813068453) | 7 |
| 330074324 | | SMP286(297947227) | 7 | 297925719 | SMP478 | SMP287(5813068453) | 7 |
| 295136450 | SMP303 | SMP286(297947227) | 7 | 2162779161 | | SMP287(5813068453) | 6 |
| 297917459 | SLP388 | SMP286(297947227) | 7 | 327255181 | SMP290 | SMP287(5813068453) | 6 |
| 5813047422 | | SMP286(297947227) | 6 | 2194845024 | | SMP287(5813068453) | 6 |
| 5813024295 | | SMP286(297947227) | 6 | 5812980467 | | SMP287(5813068453) | 6 |
| 5813011015 | SMP298 | SMP286(297947227) | 6 | 2226221475 | | SMP287(5813068453) | 6 |
| 5813011794 | | SMP286(297947227) | 6 | 5813012529 | DNd01 | SMP294(389563849) | 22 |
| 758320353 | | SMP286(297947227) | 6 | 5813020864 | DNd01 | SMP294(389563849) | 20 |
| 699752330 | | SMP286(297947227) | 6 | 419575044 | SLP208 | SMP294(389563849) | 13 |
| 613822435 | SMP103 | SMP286(297947227) | 6 | 5813046951 | pC1a | SMP294(389563849) | 13 |
| 580010810 | | SMP286(297947227) | 6 | 5813009881 | SMP485 | SMP294(389563849) | 11 |
| 548972278 | | SMP286(297947227) | 6 | 327937506 | | SMP294(389563849) | 9 |
| 546925626 | | SMP286(297947227) | 6 | 5813055822 | | SMP294(389563849) | 8 |
| 480698453 | SMP303 | SMP286(297947227) | 6 | 358286194 | SMP537 | SMP294(389563849) | 8 |
| 453794660 | SMP098_b | SMP286(297947227) | 6 | 5813014245 | PI1_b | SMP294(389563849) | 8 |
| 421054850 | SMP224 | SMP286(297947227) | 6 | 327242287 | SMP373 | SMP294(389563849) | 8 |
| 392325925 | SMP103 | SMP286(297947227) | 6 | 359027729 | PI1_b | SMP294(389563849) | 8 |
| 390344371 | SMP230 | SMP286(297947227) | 6 | 297947227 | SMP286 | SMP294(389563849) | 7 |
| 358277352 | | SMP286(297947227) | 6 | 327588225 | SMP083 | SMP294(389563849) | 7 |
| 326465302 | SMP225 | SMP286(297947227) | 6 | 358976440 | SMP219 | SMP294(389563849) | 7 |
| 327881747 | SLP421 | SMP286(297947227) | 6 | 327933208 | SMP486_f | SMP294(389563849) | 7 |

| | | | |
|---|---|---|---|
| 328377109 | | SMP286(297947227) | 6 |
| 298266797 | | SMP286(297947227) | 6 |
| 297593055 | SMP483 | SMP286(297947227) | 6 |
| 359744514 | pC1a | SMP286(328373131) | 110 |
| 453527730 | SMP161 | SMP286(328373131) | 46 |
| 480521586 | | SMP286(328373131) | 44 |
| 298262408 | | SMP286(328373131) | 41 |
| 327937506 | | SMP286(328373131) | 40 |
| 358290805 | SMP083 | SMP286(328373131) | 38 |
| 328632707 | SMP083 | SMP286(328373131) | 38 |
| 519694628 | SMP333 | SMP286(328373131) | 36 |
| 388638672 | SMP161 | SMP286(328373131) | 34 |
| 5813046951 | pC1a | SMP286(328373131) | 32 |
| 699752330 | | SMP286(328373131) | 30 |
| 486932201 | SMP083 | SMP286(328373131) | 27 |
| 548660840 | | SMP286(328373131) | 26 |
| 551427936 | | SMP286(328373131) | 24 |
| 513209805 | | SMP286(328373131) | 22 |
| 360763630 | | SMP286(328373131) | 22 |
| 328010530 | SMP454 | SMP286(328373131) | 22 |
| 327588225 | SMP083 | SMP286(328373131) | 21 |
| 947743009 | SMP594 | SMP286(328373131) | 20 |
| 610381395 | | SMP286(328373131) | 20 |
| 420454337 | | SMP286(328373131) | 20 |
| 329732855 | SMP368 | SMP286(328373131) | 20 |
| 547331291 | | SMP286(328373131) | 19 |
| 574260596 | | SMP286(328373131) | 18 |
| 518593663 | | SMP286(328373131) | 18 |
| 452849628 | | SMP286(328373131) | 18 |
| 423852404 | | SMP286(328373131) | 18 |
| 391121098 | | SMP286(328373131) | 18 |
| 515580115 | | SMP286(328373131) | 17 |
| 481190355 | | SMP286(328373131) | 17 |
| 5813010435 | SMP540 | SMP286(328373131) | 16 |
| 581055513 | | SMP286(328373131) | 16 |
| 483884737 | | SMP286(328373131) | 16 |
| 326253554 | SMP454 | SMP286(328373131) | 16 |
| 824620586 | | SMP286(328373131) | 15 |
| 703145276 | | SMP286(328373131) | 15 |
| 5813047717 | PI1_b | SMP294(389563849) | 7 |
| 5813057164 | PI1_b | SMP294(389563849) | 7 |
| 331006655 | SMP347 | SMP294(389563849) | 7 |
| 5813019567 | PI1_b | SMP294(389563849) | 6 |
| 481337148 | | SMP294(389563849) | 6 |
| 979065964 | FB6Z | SMP294(389563849) | 6 |
| 5813092319 | SMP346 | SMP297(5813071288) | 20 |
| 297243542 | SMP335 | SMP297(5813071288) | 17 |
| 417558532 | SMP421 | SMP297(5813071288) | 16 |
| 5813007108 | SMP346 | SMP297(5813071288) | 15 |
| 294432760 | SLP351 | SMP297(5813071288) | 14 |
| 327933027 | SMP168 | SMP297(5813071288) | 14 |
| 297925478 | SMP230 | SMP297(5813071288) | 13 |
| 267214250 | pC1b | SMP297(5813071288) | 13 |
| 298258399 | SMP577 | SMP297(5813071288) | 10 |
| 5813078074 | SMP585 | SMP297(5813071288) | 8 |
| 298258513 | SMP108 | SMP297(5813071288) | 8 |
| 266187342 | SMP347 | SMP297(5813071288) | 7 |
| 298599672 | SMP088 | SMP297(5813071288) | 7 |
| 5813055949 | SMP175 | SMP297(5813071288) | 7 |
| 448234999 | SMP230 | SMP297(5813071288) | 6 |
| 294436967 | PPL203 | SMP297(5813071288) | 6 |
| 295133015 | SLP414 | SMP297(5813071288) | 6 |

| | | | |
|---|---|---|---|
| 516944116 | | SMP286(328373131) | 15 |
| 513848395 | | SMP286(328373131) | 15 |
| 420010560 | | SMP286(328373131) | 15 |
| 330074324 | | SMP286(328373131) | 15 |
| 514527219 | | SMP286(328373131) | 14 |
| 451148412 | | SMP286(328373131) | 14 |
| 5813108370 | | SMP286(328373131) | 13 |
| 5813011794 | | SMP286(328373131) | 13 |
| 611413241 | | SMP286(328373131) | 13 |
| 578642656 | | SMP286(328373131) | 13 |
| 451040974 | SMP252 | SMP286(328373131) | 13 |
| 421815053 | | SMP286(328373131) | 13 |
| 360690414 | SMP162 | SMP286(328373131) | 13 |
| 391124880 | | SMP286(328373131) | 13 |
| 328377109 | | SMP286(328373131) | 13 |
| 5813010730 | | SMP286(328373131) | 12 |
| 5813011454 | | SMP286(328373131) | 12 |
| 5812981266 | | SMP286(328373131) | 12 |
| 641062535 | | SMP286(328373131) | 12 |
| 544590491 | | SMP286(328373131) | 12 |
| 544943707 | | SMP286(328373131) | 12 |
| 390438574 | | SMP286(328373131) | 12 |
| 328373067 | | SMP286(328373131) | 12 |
| 5813019568 | | SMP286(328373131) | 11 |
| 5813010533 | | SMP286(328373131) | 11 |
| 610722297 | | SMP286(328373131) | 11 |
| 516943720 | | SMP286(328373131) | 11 |
| 419665356 | | SMP286(328373131) | 11 |
| 421477870 | | SMP286(328373131) | 11 |
| 297588882 | | SMP286(328373131) | 11 |
| 5813039948 | | SMP286(328373131) | 10 |
| 5813014810 | | SMP286(328373131) | 10 |
| 1141035086 | | SMP286(328373131) | 10 |
| 978733232 | | SMP286(328373131) | 10 |
| 613822435 | SMP103 | SMP286(328373131) | 10 |
| 612777140 | | SMP286(328373131) | 10 |
| 582096833 | | SMP286(328373131) | 10 |
| 580348303 | | SMP286(328373131) | 10 |
| 581055643 | | SMP286(328373131) | 10 |

| | | | |
|---|---|---|---|
| 543566744 | SMP540 | SMP286(328373131) | 10 |
| 520272258 | SMP098_a | SMP286(328373131) | 10 |
| 482520889 | | SMP286(328373131) | 10 |
| 485511772 | PAL01 | SMP286(328373131) | 10 |
| 486604771 | | SMP286(328373131) | 10 |
| 450497356 | | SMP286(328373131) | 10 |
| 392748193 | | SMP286(328373131) | 10 |
| 421137107 | | SMP286(328373131) | 10 |
| 327587644 | SMP162 | SMP286(328373131) | 10 |
| 5813067452 | SMP509 | SMP286(328373131) | 9 |
| 5813046674 | | SMP286(328373131) | 9 |
| 5813047422 | | SMP286(328373131) | 9 |
| 5813039672 | | SMP286(328373131) | 9 |
| 733182574 | SMP162 | SMP286(328373131) | 9 |
| 609677179 | SMP098_b | SMP286(328373131) | 9 |
| 548972278 | | SMP286(328373131) | 9 |
| 518934633 | | SMP286(328373131) | 9 |
| 512216427 | | SMP286(328373131) | 9 |
| 487610557 | SMP094 | SMP286(328373131) | 9 |
| 482866008 | | SMP286(328373131) | 9 |
| 453794660 | SMP098_b | SMP286(328373131) | 9 |
| 423515621 | | SMP286(328373131) | 9 |
| 423512063 | | SMP286(328373131) | 9 |
| 422833681 | | SMP286(328373131) | 9 |
| 392424485 | | SMP286(328373131) | 9 |
| 297925628 | | SMP286(328373131) | 9 |
| 5901207157 | | SMP286(328373131) | 8 |
| 5813079417 | CL251 | SMP286(328373131) | 8 |
| 5813013619 | | SMP286(328373131) | 8 |
| 5813009651 | SMP480 | SMP286(328373131) | 8 |
| 5813002641 | | SMP286(328373131) | 8 |
| 921071226 | | SMP286(328373131) | 8 |
| 580719155 | | SMP286(328373131) | 8 |
| 548722538 | SMP237 | SMP286(328373131) | 8 |
| 543238468 | | SMP286(328373131) | 8 |
| 517578770 | SMP098_a | SMP286(328373131) | 8 |
| 517915796 | | SMP286(328373131) | 8 |
| 485918334 | | SMP286(328373131) | 8 |
| 455513316 | | SMP286(328373131) | 8 |

| | | | |
|---|---|---|---|
| 451485030 | | SMP286(328373131) | 8 |
| 425143700 | SMP097_a | SMP286(328373131) | 8 |
| 419802603 | | SMP286(328373131) | 8 |
| 360086302 | | SMP286(328373131) | 8 |
| 361105354 | | SMP286(328373131) | 8 |
| 360427510 | | SMP286(328373131) | 8 |
| 329728593 | | SMP286(328373131) | 8 |
| 298266950 | | SMP286(328373131) | 8 |
| 297929815 | SMP480 | SMP286(328373131) | 8 |
| 204962969 | | SMP286(328373131) | 8 |
| 5813060687 | | SMP286(328373131) | 7 |
| 5813046657 | | SMP286(328373131) | 7 |
| 5813009950 | | SMP286(328373131) | 7 |
| 5813009554 | SMP107 | SMP286(328373131) | 7 |
| 704816317 | | SMP286(328373131) | 7 |
| 674501939 | | SMP286(328373131) | 7 |
| 550983795 | | SMP286(328373131) | 7 |
| 515938422 | | SMP286(328373131) | 7 |
| 543580198 | | SMP286(328373131) | 7 |
| 486255488 | | SMP286(328373131) | 7 |
| 327596481 | SMP035 | SMP286(328373131) | 7 |
| 298586567 | | SMP286(328373131) | 7 |
| 5813111271 | SMP482 | SMP286(328373131) | 6 |
| 5813092990 | SMP107 | SMP286(328373131) | 6 |
| 5813069630 | NPFL1-I | SMP286(328373131) | 6 |
| 5813055817 | PAL01 | SMP286(328373131) | 6 |
| 5813025508 | | SMP286(328373131) | 6 |
| 5813013507 | | SMP286(328373131) | 6 |
| 641057518 | | SMP286(328373131) | 6 |
| 633447561 | AVLP472 | SMP286(328373131) | 6 |
| 605636564 | | SMP286(328373131) | 6 |
| 612073194 | | SMP286(328373131) | 6 |
| 545906392 | | SMP286(328373131) | 6 |
| 513464860 | | SMP286(328373131) | 6 |
| 519017242 | | SMP286(328373131) | 6 |
| 483539538 | | SMP286(328373131) | 6 |
| 483879756 | | SMP286(328373131) | 6 |
| 480854134 | | SMP286(328373131) | 6 |
| 453868633 | | SMP286(328373131) | 6 |

| | | | |
|---|---|---|---|
| 390702220 | | SMP286(328373131) | 6 |
| 391799394 | | SMP286(328373131) | 6 |
| 358985893 | SMP526 | SMP286(328373131) | 6 |
| 328278368 | SMP486_b | SMP286(328373131) | 6 |
| 297930119 | | SMP286(328373131) | 6 |
| 297925742 | SMP480 | SMP286(328373131) | 6 |
| 297921608 | SMP333 | SMP286(328373131) | 6 |
| 5813046951 | pC1a | SMP287(5813069072) | 60 |
| 359744514 | pC1a | SMP287(5813069072) | 47 |
| 5812981862 | SAG | SMP287(5813069072) | 40 |
| 517587356 | SAG | SMP287(5813069072) | 29 |
| 267214250 | pC1b | SMP287(5813069072) | 29 |
| 297947227 | SMP286 | SMP287(5813069072) | 28 |
| 361351171 | SMP028 | SMP287(5813069072) | 24 |
| 392821837 | pC1b | SMP287(5813069072) | 22 |
| 518930220 | SMP097_b | SMP287(5813069072) | 20 |
| 423101189 | oviIN | SMP287(5813069072) | 20 |
| 267551639 | pC1c | SMP287(5813069072) | 20 |
| 581690438 | SMP097_a | SMP287(5813069072) | 19 |
| 425143700 | SMP097_a | SMP287(5813069072) | 18 |
| 327255181 | SMP290 | SMP287(5813069072) | 18 |
| 609677179 | SMP098_b | SMP287(5813069072) | 17 |
| 327933027 | SMP168 | SMP287(5813069072) | 16 |
| 5812980467 | | SMP287(5813069072) | 15 |
| 452689494 | SMP550 | SMP287(5813069072) | 14 |
| 580348303 | | SMP287(5813069072) | 13 |
| 423512063 | | SMP287(5813069072) | 13 |
| 327242169 | SMP035 | SMP287(5813069072) | 13 |
| 297917158 | SMP035 | SMP287(5813069072) | 13 |
| 485934965 | oviIN | SMP287(5813069072) | 12 |
| 392757094 | SMP551 | SMP287(5813069072) | 12 |
| 386833850 | SMP529 | SMP287(5813069072) | 12 |
| 298599271 | SMP026 | SMP287(5813069072) | 12 |
| 5813108370 | | SMP287(5813069072) | 11 |
| 5812980326 | LHPD5e1 | SMP287(5813069072) | 11 |
| 545903200 | SMP090 | SMP287(5813069072) | 11 |
| 551427936 | | SMP287(5813069072) | 10 |
| 486932201 | SMP083 | SMP287(5813069072) | 10 |
| 451722668 | | SMP287(5813069072) | 10 |

| | | | |
|---|---|---|---|
| 390702220 | | SMP287(5813069072) | 10 |
| 328274638 | SMP551 | SMP287(5813069072) | 10 |
| 330074324 | | SMP287(5813069072) | 10 |
| 327596481 | SMP035 | SMP287(5813069072) | 10 |
| 5813071078 | SMP090 | SMP287(5813069072) | 9 |
| 5813079004 | | SMP287(5813069072) | 9 |
| 5813019568 | | SMP287(5813069072) | 9 |
| 545906392 | | SMP287(5813069072) | 9 |
| 514842241 | SMP253 | SMP287(5813069072) | 9 |
| 423174813 | | SMP287(5813069072) | 9 |
| 328373131 | SMP286 | SMP287(5813069072) | 9 |
| 5813011074 | LAL007 | SMP287(5813069072) | 8 |
| 5813013636 | | SMP287(5813069072) | 8 |
| 578642656 | | SMP287(5813069072) | 8 |
| 358290805 | SMP083 | SMP287(5813069072) | 8 |
| 328632707 | SMP083 | SMP287(5813069072) | 8 |
| 5813077469 | SMP090 | SMP287(5813069072) | 7 |
| 735470305 | SMP024 | SMP287(5813069072) | 7 |
| 704816317 | | SMP287(5813069072) | 7 |
| 550319575 | pC1c | SMP287(5813069072) | 7 |
| 550029475 | | SMP287(5813069072) | 7 |
| 544602598 | | SMP287(5813069072) | 7 |
| 487623337 | | SMP287(5813069072) | 7 |
| 512782759 | SMP090 | SMP287(5813069072) | 7 |
| 423792440 | SMP097_b | SMP287(5813069072) | 7 |
| 422622063 | SMP042 | SMP287(5813069072) | 7 |
| 416876055 | SLP278 | SMP287(5813069072) | 7 |
| 358286442 | SMP466 | SMP287(5813069072) | 7 |
| 5813092319 | SMP346 | SMP287(5813069072) | 6 |
| 5813055817 | PAL01 | SMP287(5813069072) | 6 |
| 5813055773 | SMP094 | SMP287(5813069072) | 6 |
| 5813034674 | | SMP287(5813069072) | 6 |
| 5813009589 | | SMP287(5813069072) | 6 |
| 2101740317 | | SMP287(5813069072) | 6 |
| 2132434539 | | SMP287(5813069072) | 6 |
| 611728142 | | SMP287(5813069072) | 6 |
| 609247082 | SMP024 | SMP287(5813069072) | 6 |
| 520272258 | SMP098_a | SMP287(5813069072) | 6 |
| 514527219 | | SMP287(5813069072) | 6 |

| | | | |
|---|---|---|---|
| 482783862 | | SMP287(5813069072) | 6 |
| 418261858 | SMP427 | SMP287(5813069072) | 6 |
| 327945928 | | SMP287(5813069072) | 6 |
| 297925608 | SMP084 | SMP287(5813069072) | 6 |
| 5812981862 | SAG | SMP287(5813068453) | 25 |
| 5813046951 | pC1a | SMP287(5813068453) | 24 |
| 359744514 | pC1a | SMP287(5813068453) | 23 |
| 5812980467 | | SMP287(5813068453) | 18 |
| 517587356 | SAG | SMP287(5813068453) | 17 |
| 267551639 | pC1c | SMP287(5813068453) | 17 |
| 544602598 | | SMP287(5813068453) | 16 |
| 578642656 | | SMP287(5813068453) | 12 |
| 392757094 | SMP551 | SMP287(5813068453) | 12 |
| 392821837 | pC1b | SMP287(5813068453) | 12 |
| 267214250 | pC1b | SMP287(5813068453) | 11 |
| 581690438 | SMP097_a | SMP287(5813068453) | 10 |
| 423512063 | | SMP287(5813068453) | 10 |
| 361351171 | SMP028 | SMP287(5813068453) | 10 |
| 550319575 | pC1c | SMP287(5813068453) | 9 |
| 297947227 | SMP286 | SMP287(5813068453) | 9 |
| 5813046657 | | SMP287(5813068453) | 8 |
| 512782759 | SMP090 | SMP287(5813068453) | 8 |
| 5813108370 | | SMP287(5813068453) | 7 |
| 5813061262 | | SMP287(5813068453) | 7 |
| 487623337 | | SMP287(5813068453) | 7 |
| 425143700 | SMP097_a | SMP287(5813068453) | 7 |
| 360254108 | SMP334 | SMP287(5813068453) | 7 |
| 328373131 | SMP286 | SMP287(5813068453) | 7 |
| 5813019568 | | SMP287(5813068453) | 6 |
| 550029475 | | SMP287(5813068453) | 6 |
| 545903200 | SMP090 | SMP287(5813068453) | 6 |
| 422622063 | SMP042 | SMP287(5813068453) | 6 |
| 5813071289 | SLP263 | SMP294(389563849) | 42 |
| 5813035213 | SLP079 | SMP294(389563849) | 26 |
| 481337148 | | SMP294(389563849) | 24 |
| 326892728 | SLP263 | SMP294(389563849) | 24 |
| 419575044 | SLP208 | SMP294(389563849) | 22 |
| 5812980257 | SLP263 | SMP294(389563849) | 21 |
| 389303772 | SLP263 | SMP294(389563849) | 16 |

| | | | |
|---|---|---|---|
| 388284324 | SLP032 | SMP294(389563849) | 15 |
| 450302778 | SLP264 | SMP294(389563849) | 14 |
| 480391822 | SMP532 | SMP294(389563849) | 11 |
| 297851977 | LHAV3j1 | SMP294(389563849) | 11 |
| 5813021535 | SLP230 | SMP294(389563849) | 10 |
| 484350664 | SLP086 | SMP294(389563849) | 9 |
| 420956527 | LHAV6b3 | SMP294(389563849) | 9 |
| 5813020673 | SMP252 | SMP294(389563849) | 8 |
| 481989534 | LHAV2i5 | SMP294(389563849) | 7 |
| 5813010494 | LHAV6b3 | SMP294(389563849) | 6 |
| 546412678 | SLP085 | SMP294(389563849) | 6 |
| 484355342 | LHPV5b5 | SMP294(389563849) | 6 |
| 450609707 | LHAV3b11 | SMP294(389563849) | 6 |
| 449896330 | SLP264 | SMP294(389563849) | 6 |
| 358272988 | SLP263 | SMP294(389563849) | 6 |
| 326817510 | SLP207 | SMP294(389563849) | 6 |
| 296168382 | SLP347 | SMP297(5813071288) | 21 |
| 357569231 | FS4A | SMP297(5813071288) | 20 |
| 297938108 | | SMP297(5813071288) | 20 |
| 5813082948 | FS4A | SMP297(5813071288) | 17 |
| 1036550962 | SLP355 | SMP297(5813071288) | 16 |
| 357944930 | SMP539 | SMP297(5813071288) | 16 |
| 295133743 | SLP414 | SMP297(5813071288) | 16 |
| 296509673 | | SMP297(5813071288) | 16 |
| 1010731508 | FS4A | SMP297(5813071288) | 14 |
| 327566137 | FS4A | SMP297(5813071288) | 14 |
| 5813098375 | SLP347 | SMP297(5813071288) | 13 |
| 1042495376 | FS4A | SMP297(5813071288) | 12 |
| 327565511 | FS4A | SMP297(5813071288) | 12 |
| 265120589 | SLP414 | SMP297(5813071288) | 12 |
| 5901195361 | SLP268 | SMP297(5813071288) | 11 |
| 5813041131 | FS4A | SMP297(5813071288) | 11 |
| 5813009352 | SLP182_c | SMP297(5813071288) | 10 |
| 5813009260 | SLP414 | SMP297(5813071288) | 10 |
| 357923095 | SMP539 | SMP297(5813071288) | 10 |
| 357909813 | FS4A | SMP297(5813071288) | 10 |
| 327877252 | SLP268 | SMP297(5813071288) | 10 |
| 327225145 | FS4A | SMP297(5813071288) | 10 |
| 5813067326 | SLP351 | SMP297(5813071288) | 9 |

| | | | |
|---|---|---|---|
| 948704770 | FS4A | SMP297(5813071288) | 9 |
| 511349908 | SMP183 | SMP297(5813071288) | 9 |
| 388612388 | FS4A | SMP297(5813071288) | 9 |
| 388608485 | FS4A | SMP297(5813071288) | 9 |
| 357569240 | FS4A | SMP297(5813071288) | 9 |
| 264822904 | SLP414 | SMP297(5813071288) | 9 |
| 296867934 | FS4A | SMP297(5813071288) | 9 |
| 5813128313 | FS4A | SMP297(5813071288) | 8 |
| 5813069648 | LNd | SMP297(5813071288) | 8 |
| 5813039875 | SLP091 | SMP297(5813071288) | 8 |
| 1261129931 | FS4A | SMP297(5813071288) | 8 |
| 296846768 | LHPV6f3_b | SMP297(5813071288) | 8 |
| 5813039877 | SLP091 | SMP297(5813071288) | 7 |
| 5813009350 | SMP095 | SMP297(5813071288) | 7 |
| 326875404 | FS4A | SMP297(5813071288) | 7 |
| 295828029 | SMP431 | SMP297(5813071288) | 7 |
| 5813067337 | SLP346 | SMP297(5813071288) | 6 |
| 5813026589 | SMP082 | SMP297(5813071288) | 6 |
| 5812979995 | SMP433 | SMP297(5813071288) | 6 |
| 5812979938 | SLP355 | SMP297(5813071288) | 6 |
| 418875451 | LHPV6a9_b | SMP297(5813071288) | 6 |
| 298275090 | | SMP297(5813071288) | 6 |
| 295785459 | LHAV3a5 | SMP297(5813071288) | 6 |
| 298254517 | SMP082 | SMP297(5813071288) | 6 |
| 295133015 | SLP414 | SMP297(5813071288) | 6 |
| 265120467 | SLP414 | SMP297(5813071288) | 6 |
| 296855477 | | SMP297(5813071288) | 6 |

#### Table S4. Statistical analysis results

| Groups (Figure, Fly) | Sample size (NC,NH,EC,EH) | coef | exp(coef) | 95%CI (lower, upper) | P-value |
|---|---|---|---|---|---|
| 1D, Control | 43,42,45,45 | 0.7802 | 2.182 | 1.1999, 3.9679 | 0.01054976 |
| 1D, dsx∩Gad1>Gad1-RNAi | 39,38,41,41 | 0.2309 | 1.2598 | 0.6685, 2.3741 | 0.4750354 |
| 2D, Control | 40,40,37,39 | 0.871 | 2.388 | 1.112, 5.132 | 0.02568778 |
| 2D, dsx∩Gad1$^{brain}$>Gad1-RNAi | 40,40,40,40 | -0.4211 | 0.6563 | 0.3305, 1.3034 | 0.2289632 |
| 4A, Control | 40,40,37,39 | 0.871 | 2.388 | 1.112, 5.132 | 0.02568778 |
| 4A, dsx∩Gad1$^{brain}$>Rdl-RNAi | 41,43,41,40 | 0.223 | 0.8 | 0.425, 1.506 | 0.489684 |
| 4C, Control | 41,42,40,43 | 0.8315 | 2.2968 | 1.0565, 4.9931 | 0.03584471 |
| 4C, dsx∩Gad1$^{brain}$>Dop1R2-RNAi | 43,39,60,62 | -0.3585 | 0.6987 | 0.3284, 1.4868 | 0.3520828 |
| S1, CSH | 41,39,40,39 | 1.0484 | 2.8531 | 1.3577, 5.9955 | 0.005657221 |
| S2, Control | 41,42,40,43 | 0.8315 | 2.2968 | 1.0565, 4.9931 | 0.03584471 |
| S2, dsx∩Gad1$^{brain}$>Gad1-RNAi | 41,42,40,43 | -0.3918 | 0.6759 | 0.2841, 1.6078 | 0.3756019 |
| S3, Control | 42,37,46,37 | 0.4695 | 1.5991 | 0.7171, 3.5663 | 0.2512853 |
| S3, dsx∩Gad1$^{brain2}$>Gad1-RNAi | 38,36,48,44 | 0.023 | 1.0241 | 0.5926, 1.7695 | 0.9321146 |
| S8, Control | 41,42,40,43 | 0.8315 | 2.2968 | 1.0565, 4.9931 | 0.03584471 |
| S8, dsx∩Gad1$^{brain}$>Dop1R1-RNAi | 42,43,32,36 | 1.0458 | 2.8457 | 1.1296, 7.1689 | 0.02651759 |
| S8, dsx∩Gad1$^{brain}$>Dop2R-RNAi | 32,39,40,40 | 0.93509 | 2.54744 | 1.15966, 5.596 | 0.01986544 |

| Group | RMTL (min) | Standard error | adjusted P-value |
|---|---|---|---|
| NC, Control | 20.37458 | 1.46937 | 0.0765436 |
| NC, dsx∩Gad1$^{brain}$>Rdl-RNAi | 24.59106 | 1.039173 | |
| NH, Control | 18.71875 | 1.770813 | 1 |
| NH, dsx∩Gad1$^{brain}$>Rdl-RNAi | 19.66589 | 1.546134 | |
| EC, Control | 18.57117 | 1.646586 | 0.05740152 |
| EC, dsx∩Gad1$^{brain}$>Rdl-RNAi | 23.51138 | 1.166228 | |
| EH, Control | 11.22436 | 1.685797 | 0.002225616 |
| EH, dsx∩Gad1$^{brain}$>Rdl-RNAi | 19.40375 | 1.665047 | |
| NC, Control | 24.08659 | 1.228292 | 0.8986436 |
| NC, dsx∩Gad1$^{brain}$>Dop1R2-RNAi | 21.89302 | 1.324734 | |
| NH, Control | 18.70556 | 1.747245 | 0.8593916 |
| NH, dsx∩Gad1$^{brain}$>Dop1R2-RNAi | 15.57479 | 1.821614 | |
| EC, Control | 23.86333 | 1.057094 | 3.24E-11 |
| EC, dsx∩Gad1$^{brain}$>Dop1R2-RNAi | 11.48361 | 1.470172 | |
| EH, Control | 11.45039 | 1.763572 | 0.6431636 |
| EH, dsx∩Gad1$^{brain}$>Dop1R2-RNAi | 8.37957 | 1.297858 | |

#### Key Resources Table

| REAGENT or RESOURCE | SOURCE | IDENTIFIER | Additional information |
|---|---|---|---|
| **Genetic reagent** | | | |
| *Canton-S* | PMID: 24086330, Gift from Dr. K. Ito | NA | Hotta-lab strain |
| *Gad1-Gal4AD, dsx-Gal4DBD* | made in this paper | NA | RRID: BDSC_60322, (6) |
| *UAS-Gad1-TRiP-RNAi attP40* | Bloomington Drosophila Stock Center | RRID:BDSC_51794 | (7) |
| *UAS-Gad1-TRiP-RNAi attP2* | Bloomington Drosophila Stock Center | RRID:BDSC_28079 | (8) |
| *TRiP-BG attP40* | Bloomington Drosophila Stock Center | RRID:BDSC_36304 | |
| *TRiP-BG attP2* | Bloomington Drosophila Stock Center | RRID:BDSC_36303 | |
| *Otd-FLP, tubP FRT-Gal80-FRT* | Gift from Dr.Miwa | NA | RRID: BDSC_38880, (9) |
| *20XUAS-IVS-mCD8::GFP* | Bloomington Drosophila Stock Center | RRID:BDSC_32194 | |
| *UAS-DenMark* | Bloomington Drosophila Stock Center | RRID:BDSC_33064 | |
| *R71G01-LexA* | Bloomington Drosophila Stock Center | RRID:BDSC_54733 | |
| *lexAop-CD4::spGFP11; UAS-CD4::spGFP1-10* | made in this paper | NA | Gift from Kristin Scott |
| *UAS-Rdl Trip RNAi attp40* | Bloomington Drosophila Stock Center | RRID:BDSC_52903 | (10) |
| *UAS-Dop1R1 Trip RNAi attp2* | Bloomington Drosophila Stock Center | RRID:BDSC_31765 | (11) |
| *UAS-Dop1R2 Trip RNAi attp2* | Bloomington Drosophila Stock Center | RRID:BDSC_26018 | (12) |
| *UAS-Dop2R Trip RNAi attp2* | Bloomington Drosophila Stock Center | RRID:BDSC_26001 | (13) |
| *tsh-Gal80* | Gift from Julie H. Simpson | NA | |
| **Software** | | | |
| Fiji (version: 2.9.0/1.53t) | PMID: 22743772 | RRID: SCR_002285 | |
| VVDViewer (version 1.5.10) | https://github.com/takashi310/VVDViewer | RRID:SCR_021708 | |
| The Audacity Team (version 2.0) | https://www.audacityteam.org/ | RRID: SCR_00719 | |
| **Tool** | | | |
| R (version 4.1.0) | https://www.r-project.org/ | RRID: SCR_001905 | |
| Neuprint | https://neuprint.janelia.org/ | NA | |
| Virtual Fly Brain | https://www.virtualflybrain.org/ | RRID:SCR_004229 | |
| SCope | https://scope.aertslab.org/ | NA | |
| **Antibody** | | | |
| Rat anti-GFP (Rat Monoclonal) | NACARAI TESQUE, INC | RRID: AB_221569 | (1:1000) |
| Mouse anti-GFP (Mouse Monoclonal) | Sigma-Aldrich | RRID: AB_259941 | (1:300) |
| Mouse nc82-s (mouse monoclonal; supernatant) | DSHB | RRID: AB_2314866 | (1:20) |
| Rabbit anti-CD4 (rabbit polyclonal) | Sigma-Aldrich | RRID: AB_1078466 | (1:300) |
| Chicken anti-GFP (chicken polyclonal) | Abcam | RRID: AB_300798 | (1:200) |
| Goat anti-rat-Alexa 488 (goat polyclonal) | Jackson ImmunoResearch | RRID: AB_2338362 | (1:300) |

| | | | |
|---|---|---|---|
| Goat anti-mouse-Alexa 555 (goat polyclonal) | Thermo Fisher Scientific | RRID: AB_141780 | (1:300) |
| Goat anti-chicken-Alexa 488 (goat polyclonal) | Thermo Fisher Scientific | RRID: AB_2534096 | (1:300) |
| Goat anti-rabbit-Alexa 555 (goat polyclonal) | Thermo Fisher Scientific | RRID: AB_2535850 | (1:300) |
| Goat anti-mouse-Alexa 647 (goat polyclonal) | Thermo Fisher Scientific | RRID: AB_2535805 | (1:300) |
